# Playing by their own rules: clade-specific trends in mandibular form and function disrupt current paradigms of early tetrapod evolution

**DOI:** 10.64898/2026.10.15.759488

**Authors:** Catherine R. C. Strong, Amandine Gillet, Robert J. Brocklehurst, Stephanie E. Pierce

## Abstract

Quantitative research into the evolution of vertebrate feeding has largely focused on the mandible, using morphofunctional changes in the lower jaw as an indication of broader shifts in overall feeding mode. In the context of major ecomorphological transformations, such as the origin of tetrapods, the paradigm of a trade-off between ‘fast-jawed’, aquatic suction-feeders *versus* ‘strong-jawed’, terrestrial biters is particularly prominent. However, the assumption underlying this perspective—that the mandible constitutes a reliable proxy for overall feeding mechanisms—has yet to be rigorously validated. Here, we test this widespread notion by examining both morphology (via geometric morphometrics) and function (via commonly-used functional metrics) across a broad dataset of extant and extinct vertebrate mandibles. We find that extant clades exhibit phylogenetically-structured morphofunctional solutions to the challenge of feeding in different environments, distinct from the trends exhibited by extinct taxa, thus highlighting the need for caution when relying on mandibles for deep-time reconstructions of feeding mode. By providing the first large-scale validation of functional trait differences across vertebrate feeding mechanisms, our research offers the quantitative context needed to decipher broad macroevolutionary trends in vertebrate feeding, thus better informing events such as the tetrapod water-to-land transition.

## 1. Introduction

The vertebrate feeding mechanism is a complex system, requiring synergy among the bony and soft tissues of the jaws (*1–15*), palate (*1, 6, 9, 12, 16, 17*), hyobranchial apparatus (*1–3, 5–12, 17–28*), and even postcranium (*6–8, 11, 12, 17, 18, 21, 25, 27*). As a key module of the vertebrate body plan, understanding the evolution of the feeding apparatus is essential to deciphering major ecomorphological transitions throughout vertebrate history (*29*). The mandible—or lower jaw—has received particular attention as a means of insight into macroevolutionary trends, given its role as a major feeding-related structure (*2–6, 29, 30*), its relative abundance in the fossil record (*13, 29–33*), and its presumed direct relationship with feeding performance (*29, 30, 33, 34*). Quantitative analyses of mandibular form and function have become increasingly prevalent when studying the deep-time evolution of vertebrate feeding, from ancient and modern radiations of fishes (e.g., *2, 5, 6, 14, 31, 35–43*) to early tetrapods (e.g., *32, 33, 44, 45*) to mammals (e.g., *46, 47–50*) and archosaurs (e.g., *29, 34*). These quantitative studies have increasingly focused on assessing the trade-offs in mandibular form and/or function that accompany major transitions in feeding mode, recently through the lens of performance landscapes (see e.g., *34, 36, 44, 49–51*).

The origin of tetrapods some 380-million-years ago constitutes a particularly vibrant area of such research. The transformation from aquatic, finned fishes to terrestrial, limbed tetrapods required fundamental changes throughout the body plan, with vision (*52*), respiration (*45, 53–55*), hearing (*56*), and locomotion (*57, 58*) among the myriad body systems whose adaptations enabled this first foray of vertebrates onto land. Particularly critical to this ecological transition was the evolution of feeding: whereas suction-feeding is highly effective in water, the much lower density and viscosity of air preclude its use on land (*8, 9, 12, 18, 44, 45, 54, 59*). Increasingly terrestrial vertebrates must therefore employ other feeding modes, a transformation typically framed among early tetrapods as evolution from suction- to biting-based—and, concomitantly, speed- to strength-dominated (*2, 3, 5, 6, 40*)—feeding (e.g., *12, 13, 16, 33, 44, 45, 59–63*). However, when and how this transition occurred remains uncertain. Whereas analyses of overall skull anatomy generally support a shift toward biting-based feeding in the earliest tetrapods (*16, 59, 61, 63, 64*) or even tetrapodomorph fishes (*54, 60*), studies focusing on mandibular biomechanics instead typically argue that feeding modes remained unchanged until much later (*31–33, 44, 45*). Given the ubiquity of mandibles in studying the deep-time evolution of vertebrate feeding, understanding why these signals differ so dramatically is a critical question of modern vertebrate palaeontology.

The widespread reliance on mandibles to assess macroevolutionary trends in feeding ecology relies upon a key assumption: that the mandible alone is an accurate proxy for overall feeding mode, such that jaw form and/or function are entirely sufficient to determine a given animal’s feeding mechanism and how it evolved through deep time. However, despite widespread acceptance, this implicit notion bears scrutiny. Most fundamentally, the underlying paradigm of a trade-off between ‘fast-jawed suction-feeders’ *versus* ‘strong-jawed biters’ was inspired primarily by actinopterygian fishes (see e.g., *2, 3, 5, 6, 40*), but has yet to be rigorously tested within a broader phylogenetic context. The potential universality of this trade-off—both phylogenetically (across other extant clades) and evolutionarily (throughout the fossil record)—is particularly critical when considering early tetrapods, given their position in bridging the gap between fishes and tetrapods (Figure 1A). This emphasis on a binary trade-off also risks eliding the existence of other feeding mechanisms; for example, many extant vertebrates exhibit both suction generation and jaw prehension (Table S1), a ‘hybrid’ condition also proposed for taxa near the water-to-land transition such as *Tiktaalik* (*54*) (see also discussion in *17, 54,* and *61* recognizing this complexity). Finally, implicit to this paradigm is the notion of a one-to-one connection between different levels of anatomical organization; i.e., that jaw function can be determined using jaw shape, and, more broadly, that the function of a complex system (the overall feeding mechanism) can be captured by just one of its constituent components (the mandible). However, given the prevalence of one-to-many, many-to-one, and even many-to-many mapping within the feeding apparatus of many taxa (see e.g., *21, 29, 39, 49, 50, 65–68*)—as well as the role of other structural (e.g., hyobranchium; *1, 7, 12, 18, 26*) and non-structural (e.g., kinematics (*7, 18, 22, 26*); motor control (*6, 23, 69*); external stimulus response (*70*)) factors in governing feeding performance—a more thorough investigation of potential decoupling between form and function is necessary.

**Figure 1.**
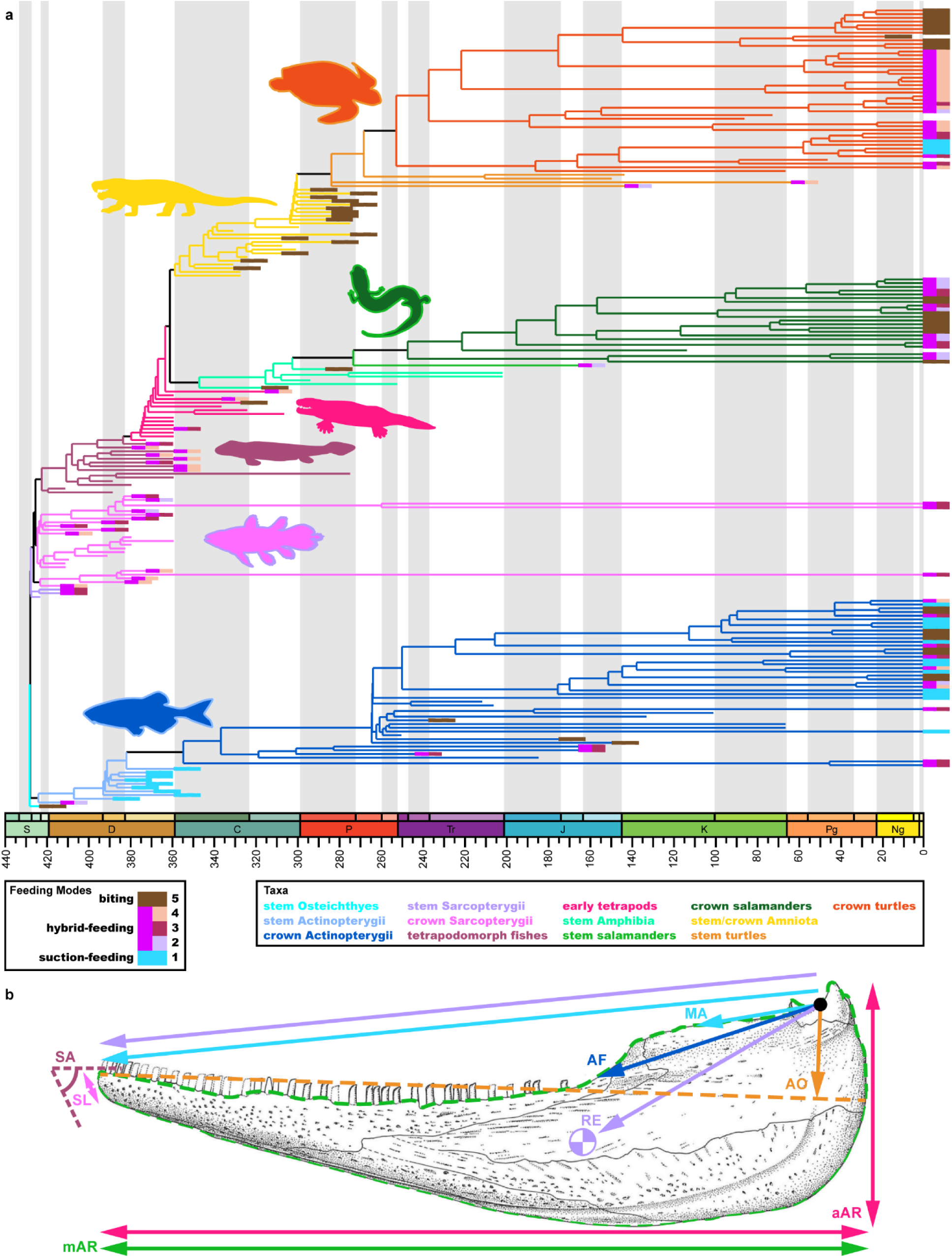
Dataset for analyzing jaw form and function. (**a**) Taxon sample and phylogenetic context. Branches colour-coded by higher taxon. Coloured boxes at tips indicate feeding modes, as scored into either three (left column) or five (right column) discrete categories; see Methods for details on how feeding modes were assigned. (**b**) Schematic of functional metrics. See Methods for details on how metrics were collected and calculated. Illustration of *Archeria* mandible modified from Holmes (*77*:fig. 14). All taxon illustrations available from PhyloPic under a CC0 1.0 Universal Public Domain Dedication license. See Methods for abbreviations of functional metrics.

Here, we explore vertebrate-wide trends in mandibular form and function, using a dataset of over 200 taxa spanning phylogenetically from fishes to tetrapods, temporally from Paleozoic to modern taxa, and ecomorphologically across several feeding modes and habitats (Figure 1; Table S1). We test a series of assumptions that, despite being implicit to past mandible-focused studies, have never been rigorously validated across vertebrates: first, that suction-feeding *versus* biting are functionally adapted for mandibular speed *versus* strength, respectively; second, that this functional specialization also manifests morphologically as a transition from gracile-jawed suction-feeders to robust-jawed biters; third, that a fundamental trade-off exists between these traits, such that a given organism can be optimized for either speed or strength, but not both simultaneously; and finally, that these morphofunctional trends reflect universal “rules” governing all vertebrates, from fishes through to tetrapods and fossils through to modern species. By rigorously testing these assumptions across an ecomorphologically diverse sample of extant and extinct vertebrates, our analyses fill an important conceptual and empirical gap essential to accurately evaluating trends in feeding mode throughout vertebrate evolution.

## 2. Results

### 2.1. Suction-feeding and biting are not differentially optimized for mandibular speed versus strength

Based on seminal research into fish feeding (*2, 3, 5, 6, 40*), suction-feeding *versus* biting are widely assumed to be optimized for mandibular speed *versus* strength, respectively (see e.g., *1, 23, 36, 37, 44, 51, 60, 71, 72*), a framework fundamental to large-scale, macroevolutionary research into early tetrapod feeding (see e.g., *32, 33, 44, 45*). To test whether this paradigmatic assumption holds true across vertebrates, we compiled a dataset of 214 mandibular silhouettes, encompassing major extant vertebrate groups (fishes, salamanders, and turtles) with diverse feeding ecologies (suction-feeding, biting, and hybrid-feeding), as well as relevant extinct taxa (early tetrapods and their relatives, and stem and crown members of the aforementioned groups) (Figure 1A, Table S1). Using proxies for jaw function as commonly implemented in prior studies (e.g., *14, 31, 32, 36, 44, 45, 50*) (Figure 1B), we used these silhouettes to measure functional metrics capturing jaw speed (average inverse aspect ratio [aAR], maximum inverse aspect ratio [mAR], rotational efficiency [RE]), jaw strength (mechanical advantage [MA], relative adductor fossa length [AF], relative symphyseal length [SL]), and other aspects of jaw function (relative articular offset [AO], symphyseal angle [SA]), allowing us to test whether these metrics distinguish among feeding modes as expected. To investigate functional trends and trade-offs across our vertebrate sample, we performed principal components analyses (PCA) of our functional metrics to generate “mechanospaces” for various partitions of our overall dataset (see Methods and Table S2), focusing on six key groupings: all taxa, all extant taxa, fishes, salamanders, turtles, and extinct taxa directly spanning the water-to-land transition (Figure 2).

**Figure 2.**
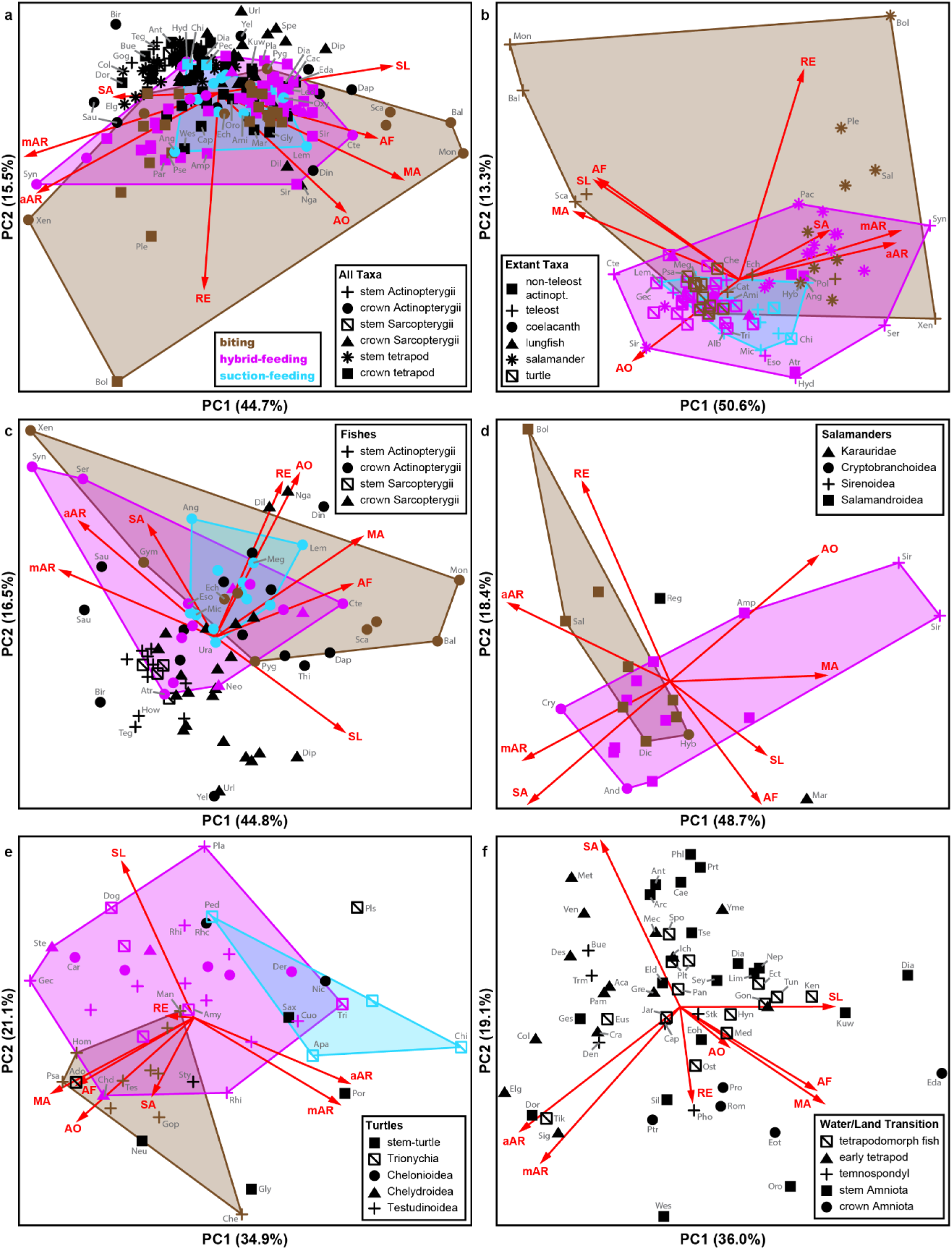
Mechanospace for key taxon partitions, reflecting principal components analysis of functional metrics. Refuting the notion of a universal trade-off between ‘rapid suction-feeders’ *versus* ‘strong biters’, major feeding modes instead exhibit similar functional proxies at a broad phylogenetic scope, with different taxon partitions following varied trends in jaw speed and strength. (**a**) All taxa (n = 214). (**b**) Extant taxa (n = 95). (**c**) Fishes (n = 84). (**d**) Salamanders (n = 24). (**e**) Turtles (n = 48). (**f**) Taxa spanning the tetrapod water-to-land transition (n = 58). Colours indicate feeding modes for extant taxa; see legend in (a) for colour scheme used throughout all panels. Black datapoints denote extinct taxa. Shape denotes higher taxon; see respective legend for each panel. Biplots indicate the respective contribution of each functional metric to the first two principal components. See Methods for abbreviations of functional metrics and taxa.

Our analyses reveal that suction-feeders and biters do not differentially emphasize speed *versus* strength, respectively, and that feeding mode (suction, hybrid, biting) cannot be reliably predicted from these mandibular functional proxies. At a vertebrate-wide scale (Figure 2A–B), strength- and speed-related metrics are generally oriented in different directions in mechanospace (except for RE, which lies intermediate to these extremes); however, these properties are not aligned with suction-feeding *versus* biting. Instead, all feeding modes extend along this continuum, with suction-feeders falling entirely within the region occupied by hybrid-feeders (all taxa: *F*_1,65_ = 0.178, *p* = 0.787; extant taxa: *F*_1,65_ = 0.386, *p* = 0.599) and biters overlapping both suction-feeders (all taxa: *F*_1,42_ = 1.174, *p* = 0.294; extant taxa: *F*_1,42_ = 2.060, *p* = 0.131) and hybrid-feeders (all taxa: *F*_1,77_ = 1.375, *p* = 0.241; extant taxa: *F*_1,77_ = 2.322, *p* = 0.109) (Figure 2A–B). The most notable exploration of mechanospace instead occurs among biters; this feeding mechanism spans from specialized ‘rasping’ taxa (e.g., parrotfishes and triggerfish), which emphasize strength-related traits (symphyseal length, adductor fossa length, and mechanical advantage), to taxa with elongate and gracile jaws (e.g., needlefish, some salamanders), which emphasize speed-related traits (rotational efficiency, maximum inverse AR, average inverse AR) (Figure 2A–B). Size is not a major driver of this distribution across mechanospace, with regressions of PCs 1–3 against jaw size (see Methods) typically being either non-significant (PGLS: *p* > 0.05 for PC1 of the all-taxa and extant-taxa mechanospaces) and/or of very low explanatory value (R^2^ < 0.1 for all axes except PC2 of the all-taxa and extant-taxa mechanospaces) (Table S3).

Analyses of individual taxon partitions clarify this lack of vertebrate-wide differentiation among feeding modes, revealing that functional trends manifest differently across different taxa. For example, fishes follow a similar trend as described above, with all feeding modes containing a mixture of both ‘fast-’ or ‘strong-jawed’ taxa (Figure 2C). In contrast, turtles exhibit the expected trend of biters tending to emphasize strength-related traits (MA and adductor fossa length), suction-feeders tending to emphasize speed-related traits (maximum and average inverse AR), and hybrid-feeders lying in between (Figure 2E). Salamanders, though, follow an opposite relationship, with biters emphasizing speed-related traits whereas hybrid-feeders emphasize strength (Figure 2D). These analyses together indicate that suction-feeding and biting are not universally associated with functional proxies for mandibular speed *versus* strength, respectively.

To further assess the utility of mandibular functional proxies in predicting feeding modes across vertebrates, we also conducted linear discriminant analyses (LDA) using various training datasets comprising taxa of known feeding mode (Figures 3–4; Tables S4–5; see Methods for details). Regardless of which taxon partition is used to train the discriminant model, known feeding modes overlap heavily in each discriminant space (Figure 3), rather than being ‘forced’ apart as would be expected if they exhibited different functional traits. This inability to distinguish among feeding modes is further reflected in the relatively low predictive accuracy of each model, ranging from 63% for the model trained on all taxa of known feeding mode to 41% for the model trained on extant salamanders (Table 1; Figures 3–4). Underlying these numerous misclassifications, the associated posterior probabilities for each prediction are often weighted toward multiple feeding modes, rather than heavily favouring the ‘correct’ mode for that taxon as would be expected for a reliable model (Figure 4). Of particular note, these misclassifications consistently affect even commonly-accepted exemplars of each feeding mode, such as *Megalops* and *Micropterus* (suction-feeding fishes which are often misclassified as either biters or hybrid-feeders) and early-evolving amniotes (e.g., diadectomorphs and captorhinids, terrestrial herbivores which tend to be misclassified as hybrid-feeders). These discriminant models further struggle to delineate feeding modes in fossils, with the vast majority of extinct taxa (∼100 out of 119 total fossils) being predicted as ‘hybrid-feeders’ under the all-taxa-based, extant-based, and extant-fish-based models. The extant-salamander- and extant-turtle-based models do classify fossils more evenly among the possible feeding modes, but still exhibit similar misclassifications of key taxa as noted above and, in the case of salamanders, fail to predict any taxa as ‘suction-feeders’ given the absence of this feeding mode within the training dataset.

**Figure 3.**
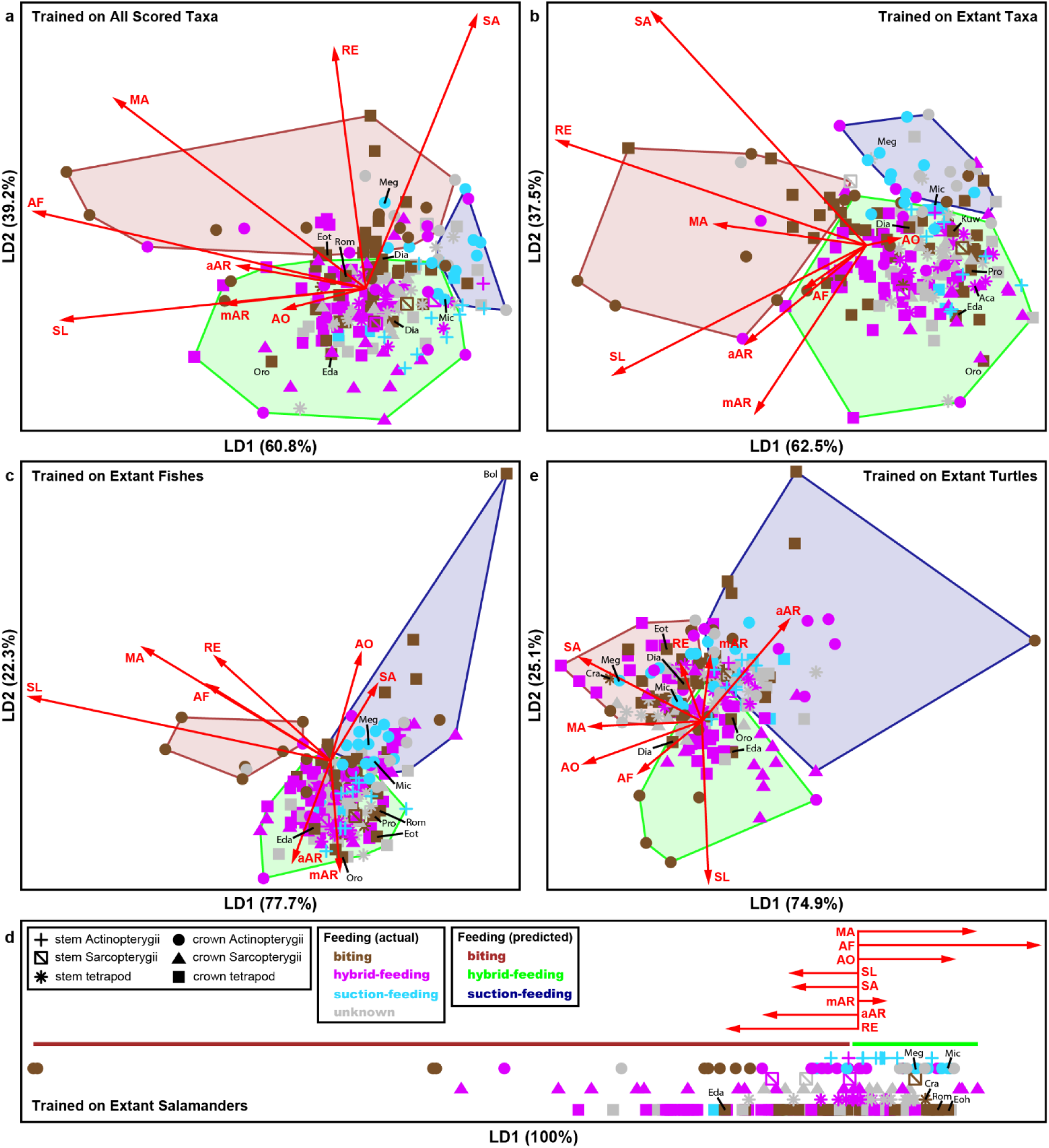
Multivariate space of key taxon partitions, reflecting linear discriminant analysis (LDA) of functional metrics. Known feeding modes overlap heavily in each discriminant space and are often mis-classified, rather than being ‘forced’ apart into accurately-assigned groups as would be expected if they exhibited different functional traits. (**a**) LDA model trained using all taxa for which feeding modes could be ascertained (n = 160, comprising 95 extant and 65 extinct taxa). (**b**) LDA model trained using all extant taxa (n = 95). (**c**) LDA model trained using extant fishes (n = 34). (**d**) LDA model trained using extant salamanders (n = 22). (**e**) LDA model trained using extant turtles (n = 39). Coloured datapoints denote known feeding modes (including among fossils), with grey representing extinct taxa for which feeding mode could not be ascertained from the literature. Coloured hulls reflect feeding modes as predicted by the discriminant model. Shape denotes higher taxon. See legends in (d) for colour schemes and shapes used throughout all panels. Biplots indicate the respective loading of each functional metric in the discriminant model. Datapoints in (d) staggered vertically for clarity. See Methods for abbreviations of functional metrics and taxa.

**Figure 4.**
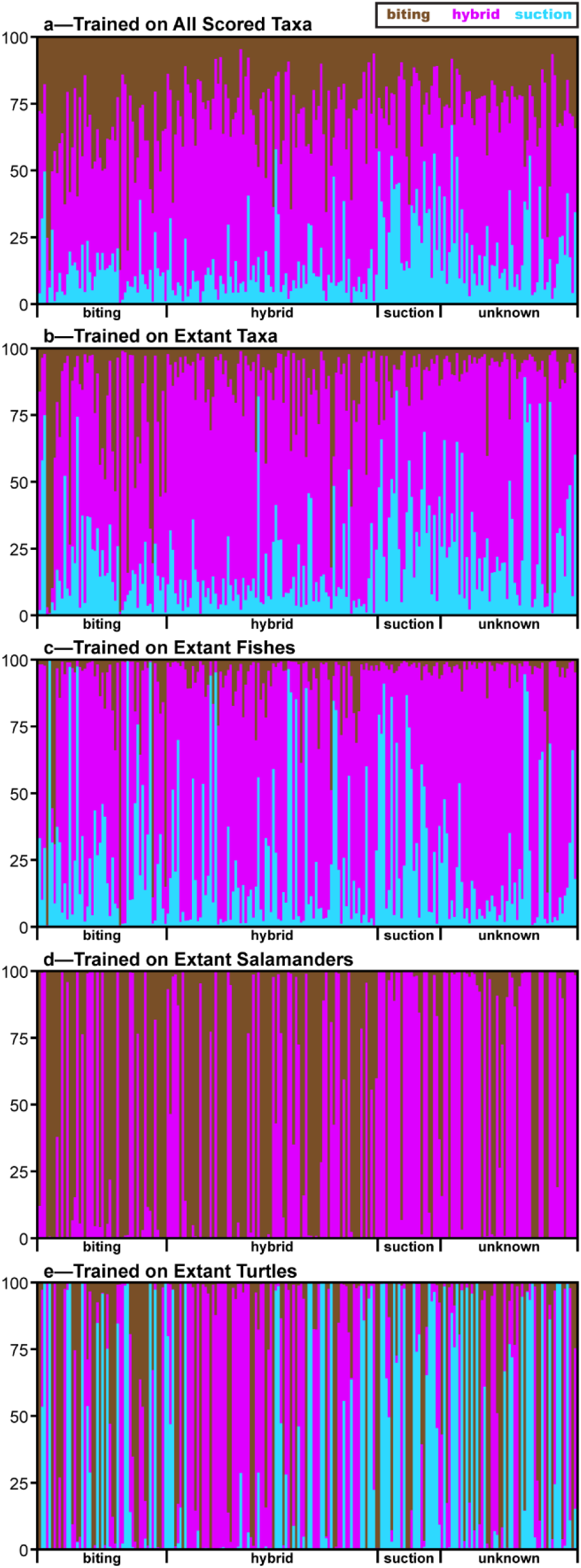
Posterior probability of feeding mode classifications, based on linear discriminant analysis (LDA) of key taxon partitions. The posterior probabilities for a given taxon are often weighted toward multiple feeding modes, rather than heavily favouring the ‘correct’ mode for that taxon as would be expected for a reliable model. (**a**) LDA model trained using all taxa for which feeding modes could be ascertained (n = 160, comprising 95 extant and 65 extinct taxa). (**b**) LDA model trained using all extant taxa (n = 95). (**c**) LDA model trained using extant fishes (n = 34). (**d**) LDA model trained using extant salamanders (n = 22). (**e**) LDA model trained using extant turtles (n = 39). Coloured bars represent posterior probability of each taxon belonging to the corresponding feeding mode; see legend at top for colour scheme used throughout all panels.

**Table 1.** Accuracy of feeding mode classifications obtained from linear discriminant analysis of key taxon partitions.

|  |  |  | *Predicted feeding mode* |  |  |
| --- | --- | --- | --- | --- | --- |
|  |  |  | Biting | Hybrid-feeding | Suction-feeding |
| *True feeding mode* | **ALL TAXA** | Biting | 19 | 30 | 2 |
|  |  | Hybrid-feeding | 10 | 72 | 2 |
|  |  | Suction-feeding | 3 | 13 | 9 |
|  |  | Unknown | 5 | 41 | 8 |
|  | **EXTANT TAXA** | Biting | 15 | 32 | 4 |
|  |  | Hybrid-feeding | 3 | 76 | 5 |
|  |  | Suction-feeding | 0 | 17 | 8 |
|  |  | Unknown | 2 | 42 | 10 |
|  | **EXTANT FISHES** | Biting | 6 | 36 | 9 |
|  |  | Hybrid-feeding | 1 | 67 | 16 |
|  |  | Suction-feeding | 0 | 15 | 10 |
|  |  | Unknown | 1 | 46 | 7 |
|  | **EXTANT SALAMANDERS** | Biting | 34 | 17 | — |
|  |  | Hybrid-feeding | 52 | 32 | — |
|  |  | Suction-feeding | 6 | 19 | — |
|  |  | Unknown | 17 | 37 | — |
|  | **EXTANT TURTLES** | Biting | 22 | 13 | 16 |
|  |  | Hybrid-feeding | 21 | 46 | 17 |
|  |  | Suction-feeding | 7 | 3 | 15 |
|  |  | Unknown | 20 | 13 | 21 |

Altogether, these analyses highlight that suction-feeders *versus* biters do not exclusively optimize speed-*versus* strength-related traits, with trends instead manifesting differently across major taxon partitions and with a high degree of functional overlap at a vertebrate-wide scope. As a result, jaw function alone—as captured by these commonly-used metrics—is therefore insufficient to distinguish feeding modes across vertebrates (Figures 2A–B and 3A–B).

### 2.2. Suction-feeders and biters are not differentially optimized for mandibular gracility versus robustness

Just as different feeding modes are often equated with different mandibular functions, so too are they typically assumed to exhibit distinct jaw shapes, with suction-feeders and biters being associated with gracile *versus* robust jaws, respectively (see e.g., *1, 23, 60*). To test whether mandibular morphology differs across feeding modes, we followed other recent studies (e.g., *34, 35, 36, 44, 45, 50, 51*) in using elliptical Fourier analysis (EFA) to quantify the two-dimensional silhouettes of the 214 mandibles in our dataset. We then performed principal components analyses to visualize trends in morphospace (Figure 5), as well as spectral clustering analyses of the resulting PC coordinates to delineate groups of taxa with statistically similar jaw shapes (Figures 6–7; Table S6).

**Figure 5.**
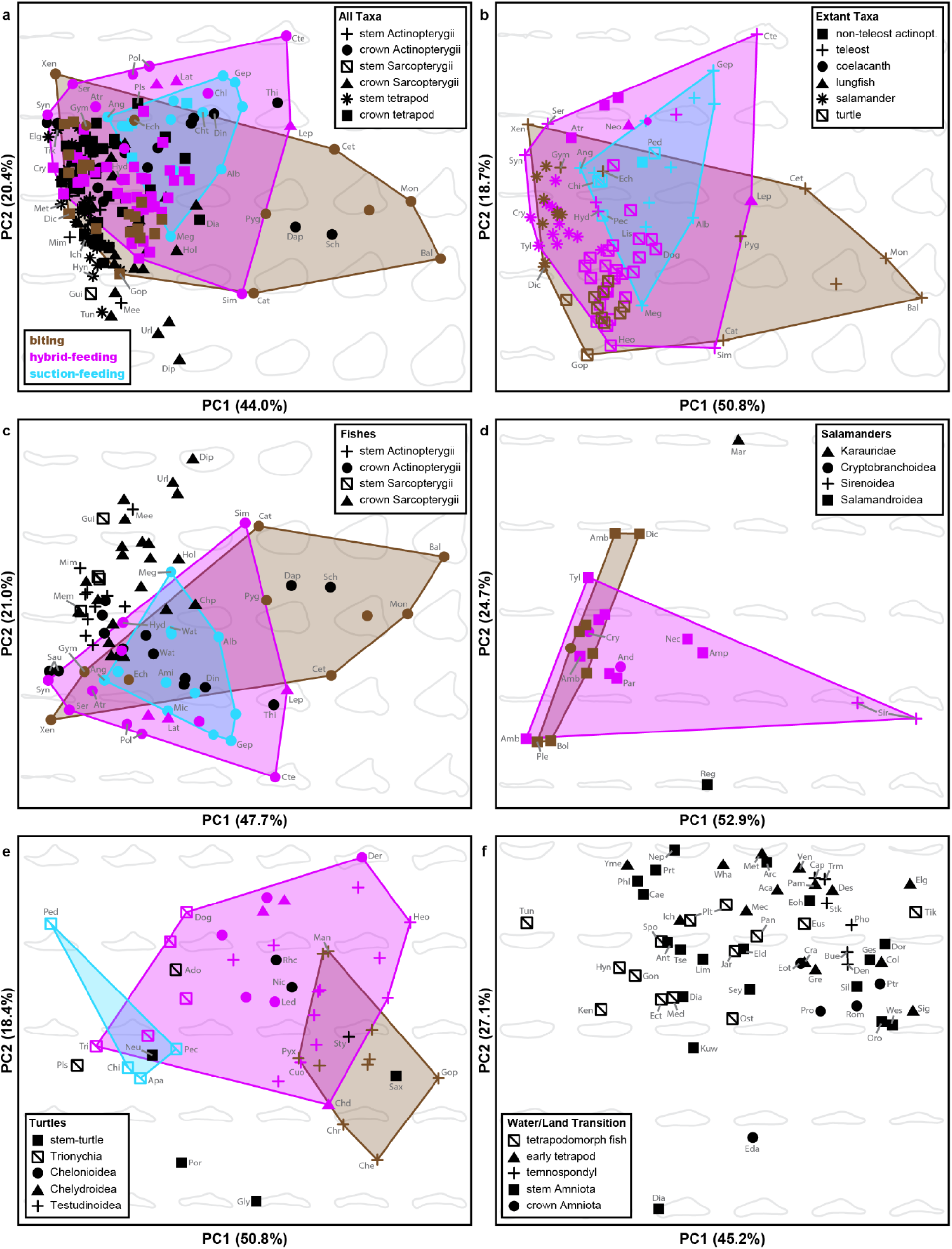
Morphospace for key taxon partitions, reflecting principal components analysis of mandible silhouettes. Refuting the notion of a universal distinction between ‘gracile suction-feeders’ and ‘robust biters’, major feeding modes instead overlap notably in jaw shape at a broad phylogenetic scope, with each taxon partition manifesting unique trends in jaw morphology. (**a**) All taxa (n = 214). (**b**) Extant taxa (n = 95). (**c**) Fishes (n = 84). (**d**) Salamanders (n = 24). (**e**) Turtles (n = 48). (**f**) Taxa spanning the tetrapod water-to-land transition (n = 58). Colours indicate feeding modes for extant taxa; see legend in (a) for colour scheme used throughout all panels. Black datapoints denote extinct taxa. Shape denotes higher taxon; see respective legend for each panel. See Methods for taxon abbreviations.

**Figure 6.**
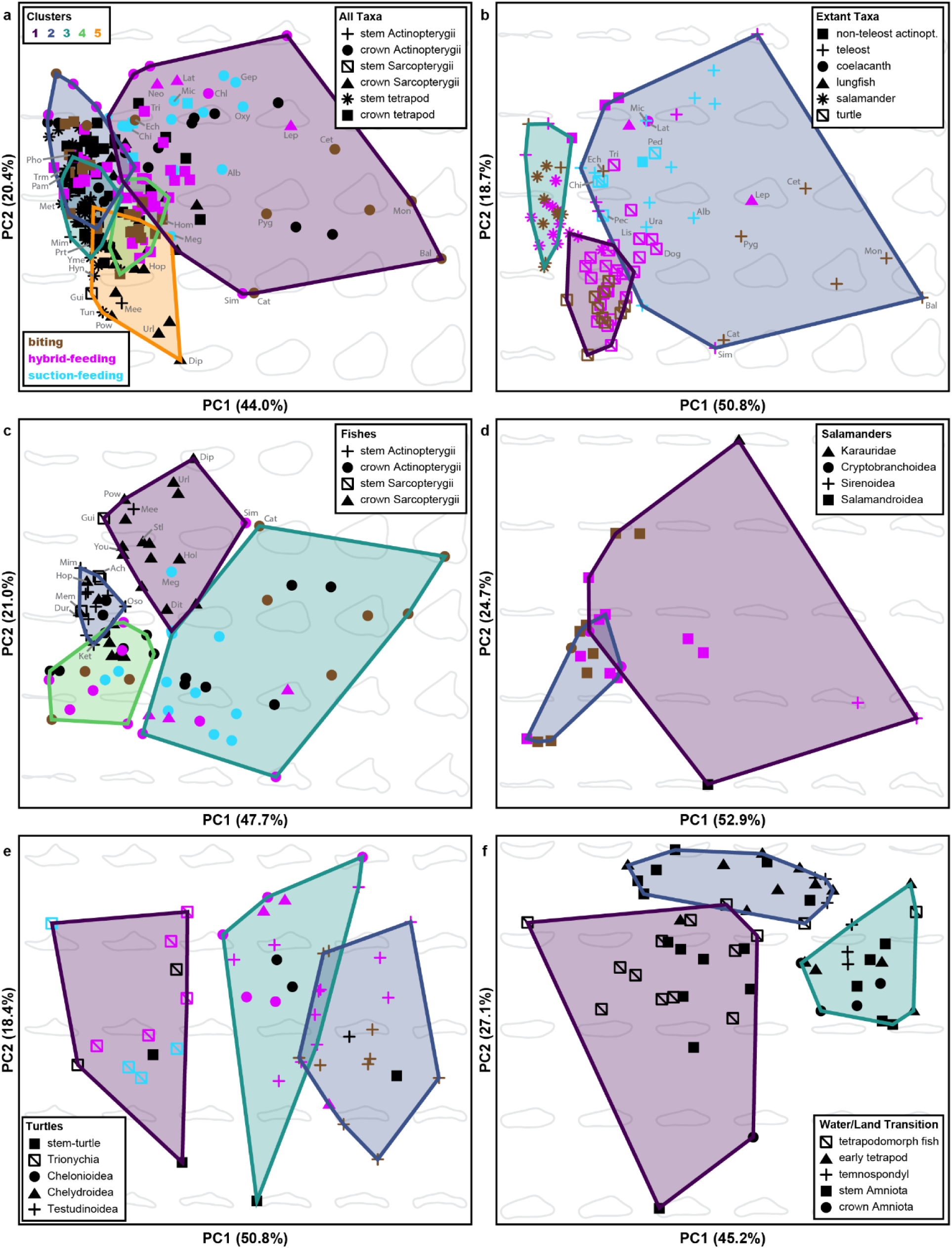
Spectral clustering analysis of key taxon partitions, as distributed throughout morphospace. Multiple feeding modes are typically recovered in the same clusters and, conversely, each feeding mode is spread across multiple clusters, indicating a lack of feeding-related signal in mandibular shape. (**a**) All taxa (n = 214). (**b**) Extant taxa (n = 95). (**c**) Fishes (n = 84). (**d**) Salamanders (n = 24). (**e**) Turtles (n = 48). (**f**) Taxa spanning the tetrapod water-to-land transition (n = 58). Coloured polygons represent cluster assignments based on PC axes >5%. Coloured datapoints indicate feeding modes for extant taxa; see legend in (a) for colour scheme used throughout all panels. Black datapoints denote extinct taxa. Shape denotes higher taxon; see respective legend for each panel. See Methods for taxon abbreviations.

**Figure 7.**
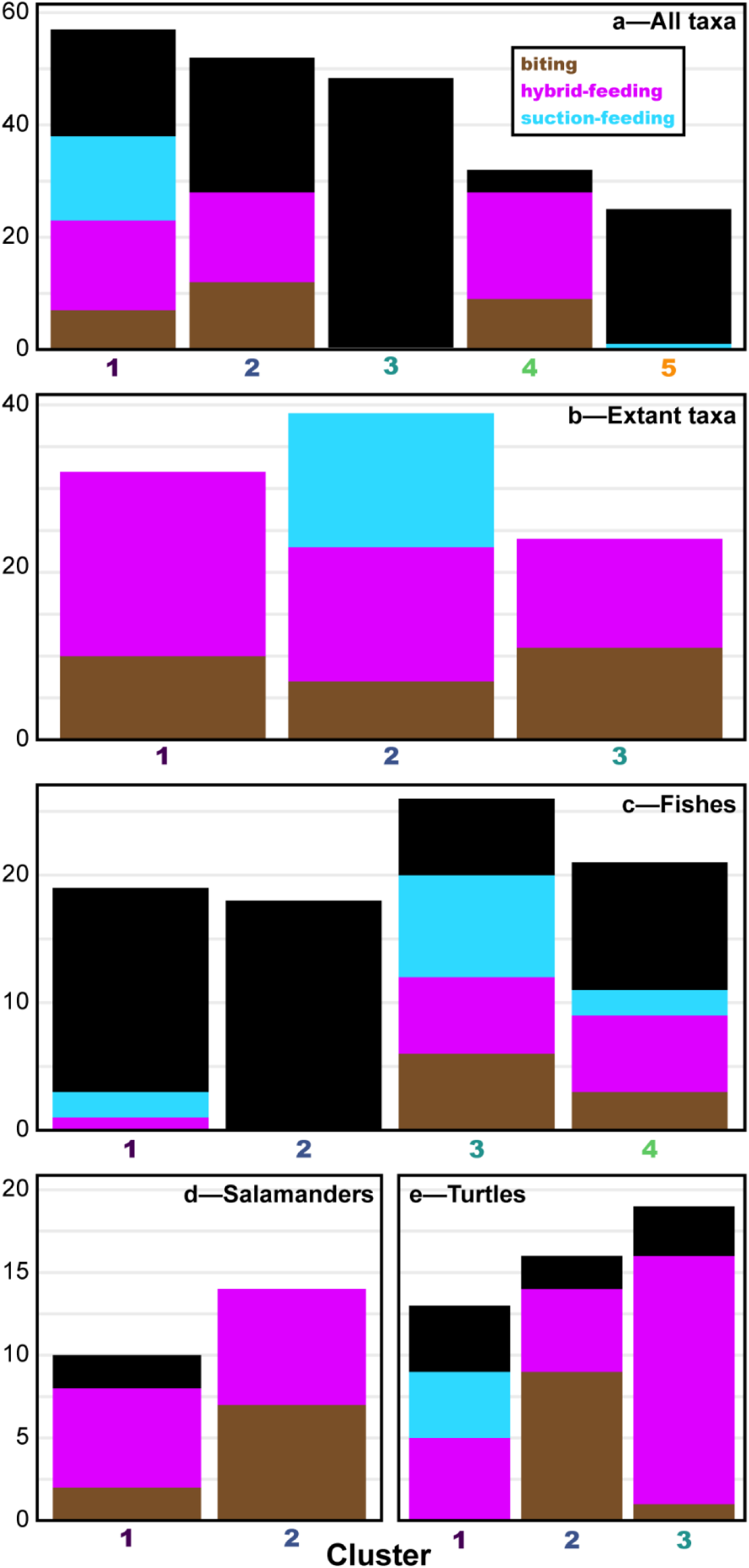
Distribution of feeding modes across morphological clusters, obtained via spectral clustering analysis of key taxon partitions. Rather than separating by feeding mode, jaw shape moreso exhibits temporal partitioning, with extinct (especially Paleozoic) taxa largely clustering with each other and separately from extant vertebrates. (a) All taxa (n = 214). (b) Extant taxa (n = 95). (**c**) Fishes (n = 84). (**d**) Salamanders (n = 24). (**e**) Turtles (n = 48). Coloured bars indicate feeding modes for extant taxa; see legend in (a) for colour scheme used throughout all panels. Black denotes extinct taxa. Coloured labels along *x*-axis correspond to cluster polygons in Figure 6.

These analyses refute the notion of a universal distinction between ‘gracile suction-feeders’ and ‘robust biters’. Although many taxon partitions—namely, all taxa (Figure 5A), extant taxa (Figure 5B), fishes (Figure 5C), and salamanders (Figure 5D)—exhibit a transition from ‘gracile’ to ‘robust’ jaws across morphospace, this axis of variation does not consistently correspond to different feeding modes. Instead, at a vertebrate-wide scale (i.e., incorporating all taxa or all extant taxa; Figure 5A–B), suction-feeders, biters, and hybrid-feeders span across this morphological spectrum, echoing the signal from our functional data (Figure 2) in showing a high degree of overlap (all taxa—B *vs* SF: *F*_1,42_ = 2.772, *p* = 0.088; B *vs* H: *F*_1,77_ = 1.859, *p* = 0.156; H *vs* SF: *F*_1,65_ = 7.285, *p* = 0.001; extant taxa—B *vs* SF: *F*_1,42_ = 3.132, *p* = 0.060; B *vs* H: *F*_1,77_ = 1.660, *p* = 0.194; H *vs* SF: *F*_1,65_ = 9.167, *p* = 0.0003). Biting fishes are again the only taxa to diversify notably from other feeding modes, diverging into two groups—stout-jawed durophages such as parrotfishes *versus* slender-jawed raptorial biters such as moray eels and needlefish—specialized on the extrema of this continuum, with stout-jawed raptorial-biting piranhas occupying a more intermediate position (Figure 5A–B).

Spectral clustering analyses of these vertebrate-wide morphospaces reinforce the lack of feeding-related signal in mandibular shape, with multiple feeding modes being recovered in the same clusters and, conversely, each feeding mode typically being spread across multiple clusters (Figures 6A–B and 7A–B). Even in instances where all representatives of a given feeding mode do occur within the same cluster—e.g., all suction-feeders from the extant morphospace being recovered in Cluster 2 (Figures 6B and 7B) or almost all suction-feeders from the all-taxa morphospace being recovered in Cluster 1 (Figures 6A and 7A)—this signal is unreliable, as these clusters also contain specialized biters (e.g., parrotfishes, triggerfish, piranhas, moray eel). These patterns of morphospace occupation again show little allometric signal, with regressions of PCs 1–3 against jaw size typically being either non-significant (PGLS: *p* > 0.05 for PCs 1 and 3 of the all-taxa morphospace, and PCs 1–2 of the extant-taxa morphospace) and/or of very low explanatory value (R^2^ ≤ 0.12 for all axes except PC3 of the all-taxa morphospace) (Table S3). Altogether, these analyses demonstrate that feeding modes do not conform to expected morphological trends, instead often exhibiting very little morphological separation on a vertebrate-wide scale.

Rather than being driven by feeding mode, these vertebrate-wide trends in jaw shape largely reflect phylogenetic history, with fishes, salamanders, and turtles each occupying distinct regions of morphospace (*F*_2,92_ = 25.387, *p* < 0.0001) and with the encroachment of softshell turtles into the ‘fish’ region being the only notable overlap among these groups (Figures 5B and 6B, Figure S1). In turn, feeding modes *within* each taxon partition do tend to exhibit greater separation when analyzed separately (Figure 5C–E), particularly among turtles and salamanders; however, the specific trends exhibited by each group are not consistent, and cluster analyses of these respective taxon partitions again reflect a lack of clear connection between cluster assignment and feeding mode (Figures 6C–E and 7C–E). For example, fishes (Figure 5C) show a similar amount of overlap among feeding modes as in the all-taxa and extant-taxa morphospaces (Figure 5A–B), with stout-jawed biters exhibiting the most notable exploration of morphospace whereas slender-jawed biters (e.g., eels, needlefish) overlap with hybrid- and suction-feeders (B *vs* SF: *F*_1,19_ = 3.430, *p* = 0.068; B *vs* H: *F*_1,20_ = 4.832, *p* = 0.030; H *vs* SF: *F*_1,23_ = 0.694, *p* = 0.505). In contrast, salamanders also span from ‘gracile’ to ‘robust’ forms, but in the opposite polarity as expected, with biters constrained to more slender morphologies whereas hybrid-feeders explore more robust forms with more prominent coronoid processes (*F*_1,20_ = 2.537, *p* = 0.098) (Figure 5D). Finally, turtles exhibit the clearest gradation across morphospace from suction-feeders to hybrid-feeders to biters (B *vs* SF: *F*_1,12_ = 35.318, *p* = 0.001; B *vs* H: *F*_1,33_ = 9.153, *p* = 0.001; H *vs* SF: *F*_1,27_ = 10.646, *p* < 0.001); however, these feeding modes differ moreso in terms of overall mandibular profile, reflecting changes to the mandibular symphysis (slender, pointed, and tilted in suction-feeders *versus* blunt and roughly vertical in biters) and coronoid process (tall and pointed in suction-feeders *versus* short and rounded in biters) (Figure 5E). Overall, the unique morphological solutions exhibited by each taxon partition therefore refute the assumption that different feeding modes are universally optimized for different mandibular shapes.

### 2.3. Mandibular speed and strength are not mutually exclusive

Intrinsic to the prevailing perspective of a mandibular speed-*versus*-strength (or, gracility-*versus*-robustness) trade-off between different feeding modes is the underlying principle that these properties are mutually exclusive, i.e., that a given organism can be optimized for either strength or speed but not both (see e.g., *30, 34, 36, 42, 44, 51, 67, 71, 73*). We directly tested this trade-off from a functional perspective by calculating the pairwise correlation among all metrics for various partitions of our dataset (Figure 8). A broad trade-off between strength and speed loosely exists: strength-related metrics are typically positively correlated among each other, as are speed-related metrics, and a negative correlation generally exists between these sets of metrics (Figure 8). However, there are many exceptions to this signal. For example, rotational efficiency rarely manifests as expected, instead showing only a weakly positive (Figure 8A,K–L)—or even negative (Figure 8C–F,H–J,L)—correlation with other speed-related metrics in most taxon partitions and, conversely, a weakly negative (Figure 8A–C,J–L) to outright positive (Figure 8A,C–F,I–L) correlation with strength-related metrics. Extant partitions (Figure 8E,H,K) further show different patterns of correlation among key traits—such as symphyseal length *versus* adductor fossa length, symphyseal length *versus* mechanical advantage, or rotational efficiency *versus* any of the speed-or strength-related traits—indicating different strength/speed-related proxy trade-offs across vertebrates.

**Figure 8.**
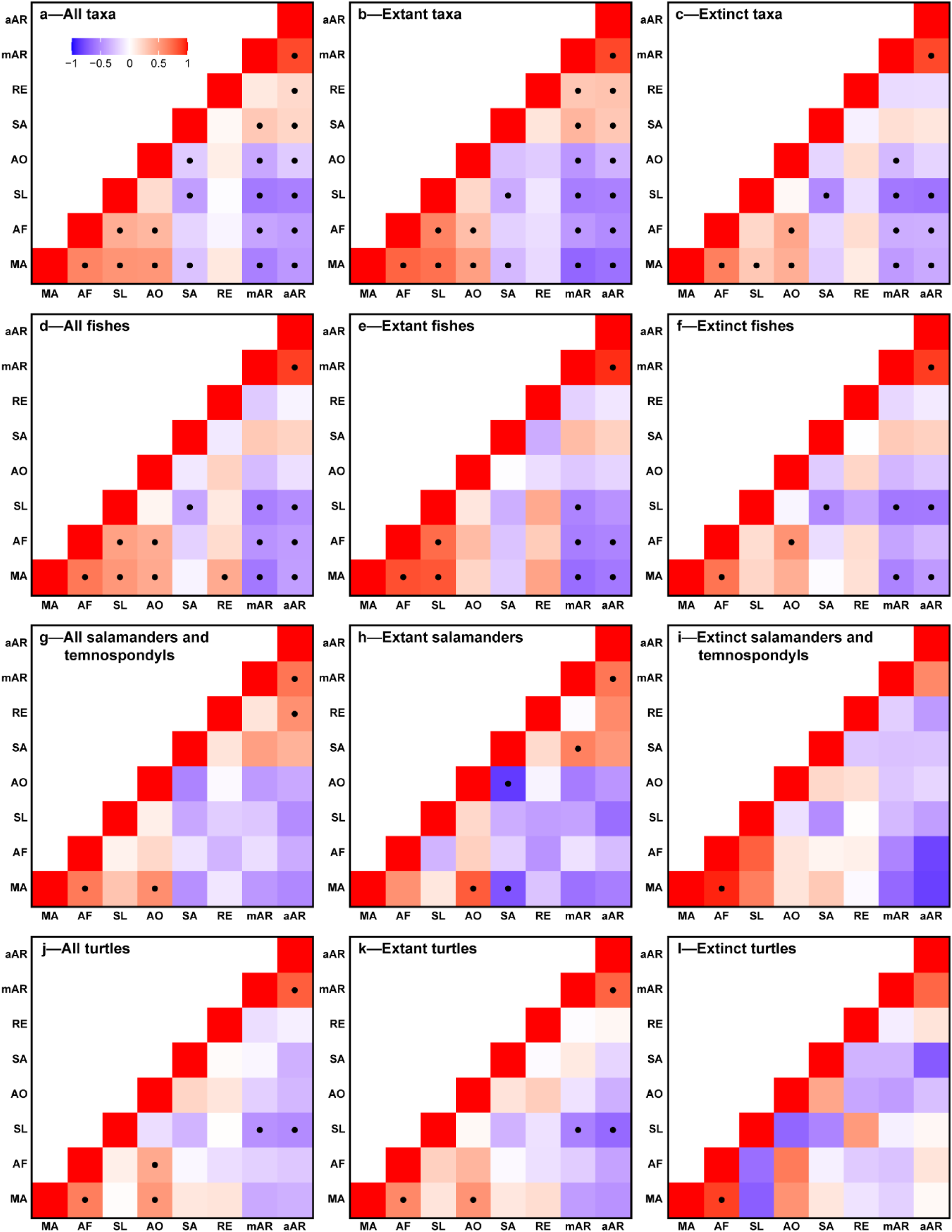
Heatmaps depicting pairwise correlation among functional metrics for key taxon partitions. Although a loose trade-off between strength and speed exists—with strength-related metrics typically being positively correlated among each other, as are speed-related metrics, and a negative correlation generally existing between them—there are many exceptions to this signal. (**a**) All taxa (n = 214). (**b**) Extant taxa (n = 95). (**c**) Extinct taxa (n = 119). (**d**) All fishes (n = 84). (**e**) Extant fishes (n = 34). (**f**) Extinct fishes (n = 50). (**g**) All salamanders and temnospondyls (n = 30). (**h**) Extant salamanders (n = 22). (**i**) Extinct salamanders and temnospondyls (n = 8). (**j**) All turtles (n = 48). (**k**) Extant turtles (n = 39). (**l**) Extinct turtles (n = 9). See scale bar in (a) for colour scale used throughout all panels. Cells marked with a dot are significant at p<0.05. See Methods for abbreviations of functional metrics.

The loadings of functional metrics in our mechanospace and discriminant-space analyses also bear on this question of functional trade-offs. Although speed- and strength-related traits do tend to be oriented toward generally different regions of mechanospace (Figure 2), these metrics are not strictly opposite to each other as would be expected under a true trade-off, indicating that an emphasis on strength-related proxies does not necessarily entail a proportional decrease in speed-related proxies and vice versa. The inconsistent relationships between mandibular functional metrics and feeding mode further amplify these trends in linear-discriminant-space: because mandibular speed *versus* strength are generally aligned with different feeding modes among salamanders and turtles, the discriminant models based on extant salamanders (Figure 3D) and on extant turtles (Figure 3E) reinforce the differing orientation of these traits from the corresponding mechanospaces (Figure 2D–E); in contrast, because mandibular function shows no connection with feeding mode in our other taxon partitions, the discriminant models based on all taxa (Figure 3A), all extant taxa (Figure 3B), and extant fishes (Figure 3C) generally collapse strength-and speed-related metrics into similar orientations (e.g., the predicted ‘biting’ region of the all-taxa discriminant model exhibits both elevated rotational efficiency and mechanical advantage). These results again indicate a complex relationship between mandibular strength and speed proxies across vertebrates.

Finally, from a morphological perspective, a spectrum from gracility to robustness does tend to be a major trend across morphospace (see Figure 5A–D), but not for all taxa, with turtles being a major exception (see Figure 5E). Altogether, although mandibular speed and strength metrics do tend to be negatively related, they are not mutually exclusive as required to constitute a strict trade-off, with our data instead revealing a more nuanced relationship amongst these traits than typically appreciated.

### 2.4. Morphofunctional trends manifest differently across major vertebrate groups and throughout deep time

The overall approach of using lower jaws to quantify the evolution of vertebrate feeding ecology stems largely from findings of mandibular speed/strength trade-offs in extant actinopterygians (e.g., *2, 3, 5, 6, 40*). In extending this principle from its original temporal and phylogenetic bases (i.e., modern actinopterygians) to broader evolutionary contexts—such as early gnathostomes (e.g., *14, 31, 36, 51*) and early tetrapods (e.g., *32, 33, 44, 45*)—this approach assumes that these form/function relationships apply universally across all vertebrates. However, based on our dataset, this assumption falls short on multiple levels.

Even within the clade that inspired this paradigm—i.e., actinopterygian fishes—the relationship between feeding mode and mandibular properties is complex: a single feeding mode can be accomplished by a range of morphologies (Figure 5C) and functional traits (Figure 2C), and, conversely, different feeding modes can exhibit similar mandibular conditions. Exemplifying this latter point, the ‘fastest’ (Figure 2C) and most gracile (Figure 5C) jaws among our dataset in fact occur in hybrid-feeders (e.g., cutthroat and sawtooth eels) and biters (e.g., garfish and moray eel), violating the expectation of biting being exclusively linked to ‘strong jaws’. More broadly, as discussed above, different partitions of extant taxa exhibit different morphofunctional trends, thus refuting the assumption of a single vertebrate-wide framework linking mandibular evolution to that of the overall feeding mechanism.

Perhaps most strikingly, our analyses reveal temporal partitioning in mandibular trends, with fossils often differing notably from their extant relatives in both form and function. For example, in the all-taxa morphospace (Figure 5A), stem-groups are much more limited in mandibular shape than their corresponding crown-groups (see also Figure S1). More broadly, extinct taxa as a whole are largely restricted to a narrow range of morphologies (Figure 5A), spanning various extents of robustness (as captured by PC2) but being mostly limited along PC1 to morphologies lacking a pronounced coronoid process. These morphologies are further notable in often exploring jaw shapes not present in extant taxa. This separation occurs both within the all-taxa morphospace (*F*_1.212_ = 20.534, *p* < 0.0001) (Figure 5A) and within each taxon-partition-specific morphospace (Figures 5C–E), although among the latter it is only statistically significant among fishes (fishes: *F*_1.82_ = 16.920, *p* < 0.0001; salamanders: *F*_1.22_ = 1.532, *p* = 0.198; turtles: *F*_1.46_ = 2.094, *p* = 0.126) (Figure 5C). Cluster analyses reveal this effect to be particularly strong among Devonian taxa: within the fish morphospace, stem-actinopterygians, stem-sarcopterygians, and extinct crown-sarcopterygians are almost entirely restricted to Clusters 1 and 2 (Figures 6C and 7C; Table S6); and, within the all-taxa morphospace, Clusters 3 and 5 comprise almost exclusively Paleozoic fishes and tetrapods (Figures 6A and 7A; Table S6).

Fossils also tend to exhibit different functional traits than extant taxa, again occupying regions of mechanospace that are both relatively limited in scope and often beyond the bounds of extant taxa (all taxa: *F*_1,212_ = 23.615, *p* < 0.0001; fishes: *F*_1.82_ = 8.238, *p* =0.001; salamanders: *F*_1.22_ = 0.682, *p* = 0.407; turtles: *F*_1.46_ = 2.548, *p* = 0.082) (Figure 2A,C–E), and with stem-taxa being much less mechanically diverse than their crown counterparts (Figure S2). In more phylogenetically-broad partitions—i.e., those incorporating all taxa (Figure 2A) or all fishes (Figure 2C)—extinct taxa are further notable in that they largely occur away from the directions in which any functional metrics are loaded, reflecting low values of many or all functional traits that suggest poor mandibular performance relative to later-evolving taxa. Finally, the patterns of correlations among functional traits also differ for extinct *versus* extant species: for example, extant fishes (Figure 8E) show a stronger positive correlation among strength-related traits than extinct fishes (Figure 8F); extant salamanders (Figure 8H) differ from extinct salamanders and temnospondyls (Figure 8I) in the correlations among strength-related traits and of rotational efficiency with speed- or strength-related traits; and, extant turtles (Figure 8K) differ from extinct turtles (Figure 8L) in the correlations among strength-related traits, in rotational efficiency *versus* symphyseal length, and in overall trade-offs between strength- and speed-related traits.

Among extinct taxa, the origin of tetrapods warrants particular attention given the emphasis on mandibles in recent studies of early tetrapod evolution (e.g., *32, 33, 44, 45*). Morphologically, our analyses reveal the tetrapod water-to-land transition to have involved a shift from deep and robust jaws with a minimal coronoid process (in earlier-diverging tetrapodomorph fishes such as osteolepidids and some tristichopterids), to gracile and curved jaws lacking a distinct coronoid process (in taxa directly surrounding the water-to-land transition, i.e., elpistostegalians, early tetrapods, temnospondyls, and early-diverging stem-amniotes), to deep and robust jaws bearing a prominent coronoid process (in later-diverging stem- and crown-amniotes) (Figures 5F and 6F; Figure S1). Functional data echo this trend (Figure 2F; Figure S2): later-diverging tetrapodomorph fishes (i.e., tristichopterids and elpistostegalians), early tetrapods, temnospondyls, and early-diverging stem-amniotes are generally localized in a region of mechanospace where none of the strength- or speed-related traits are maximized, whereas taxa farther away from this transition (i.e., earlier-diverging osteolepidid fishes, and especially later-diverging stem/crown-amniotes) explore different functions, extending toward a greater emphasis on strength-related traits. These trends therefore reflect a complex evolutionary trajectory through this phase of vertebrate evolution.

Overall, our data provide a nuanced picture of the diverse morphofunctional trends manifested across vertebrate clades and throughout deep time. As a result, although it is tempting—and would certainly be convenient—to apply the same framework of a ‘fast suction-feeder *versus* strong biter’ trade-off equally across all taxa, our results instead belie a more complex reality, one shaped by varied decoupling of the mandible from the Gestalt of the overall feeding mechanism.

## 3. Discussion

The mandible is a widely used proxy for vertebrate feeding mode, reflecting the notion that this structure alone conveys the form, function, and evolution of the overall feeding mechanism. This paradigm is especially prominent among studies concerning the evolution of terrestrial feeding across the origin of tetrapods: long conceptualized as a shift from suction-feeding to biting (see e.g., *12, 13, 16, 33, 44, 45, 59–63*), this transformation is often framed in terms of trade-offs between mandibular speed *versus* strength, respectively (see e.g., *32, 44*). In this study, we tested this widespread perspective using morphofunctional data from mandibles spanning major vertebrate groups and feeding modes. Our analyses dispel critical assumptions underlying this paradigm: different feeding modes do not differ consistently in mandibular form or function, and those trends that do exist manifest differently across vertebrates. Therefore, although analyses of the lower jaw are useful in understanding how the mandible *as one component of the overall feeding mechanism* has evolved, caution should be exercised when interpreting higher levels of biological organization.

### 3.1. The role of the mandible in interpreting vertebrate feeding

The lack of correspondence between mandibular properties and overall feeding mode, as evidenced by our data, reflects both many-to-one mapping (i.e., the same feeding mechanism being accomplished via different mandibular shapes and functions; *21, 65, 66, 68*) and one-to-many mapping (i.e., a single mandibular shape or functional metric occurring in different feeding mechanisms; *21, 46*). This decoupling in part conveys the complex contribution of the mandible to vertebrate feeding. For example, although suction-feeding is typically equated with rapid jaw movement, suction generation in reality relies upon both speed *and* volume of water ingestion (*11, 74, 75*). This feeding mode can therefore be accomplished via either “high-velocity, low-volume” *or* “low-velocity, high-volume” approaches, reflecting differences in gape size, muscular force production, and cranial linkage mechanisms (*76*). Similarly, although biting is traditionally synonymized with robust, strong-but-slow jaws, slender and rapid jaws can also prove beneficial in reducing drag during lateral snapping (*17, 29, 34, 77*) or in entrapping evasive prey (*1, 5, 34, 39, 40*). Conversely, stout, strong jaws can benefit suction-feeders by enabling the mouth to be closed against incoming rushes of water (*9*) and by providing a planar circular mouth shape that maximizes suction flow fields (*39, 78*). Therefore, although mandibular speed/strength trade-offs often exist—as is generally the case within our dataset—neither suction-feeding nor biting are exclusively limited in form or function to either extreme, enabling complex mapping of jaw properties to overall feeding modes.

Beyond the mandible, feeding mechanisms also comprise various other components such as the hyobranchial apparatus (*1–3, 5–12, 17–28*), palate (*1, 6, 9, 12, 16, 17*), teeth (*38, 39, 49, 72, 79, 80*), craniomandibular musculature (*4–6, 8, 9, 12, 17, 19, 23*), lips (*12, 81*), overall head shape (*1, 8, 9, 11, 12, 28, 61, 73*), pectoral girdle and musculature (*6–8, 11, 17, 21, 25, 27*), and axial musculature (*6, 8, 11, 12, 17, 25, 27*). Transformations to any of these structures can thus allow the overall feeding mechanism to transform independently of the lower jaw (e.g., as suggested by *29*). Furthermore, increasing the number of components in a given system in turn increases the number of possible morphological pathways by which to attain a given function (see also *49, 65*). The morphofunctional complexity of vertebrate feeding mechanisms thus promotes decoupling among different levels of biological organization (*65*), while also alleviating performance trade-offs by allowing ‘detrimental’ modifications to one structure to be mitigated by compensatory modifications in others (*43, 49, 71, 74*).

This principle also explains why different vertebrate groups exhibit different trends in mandibular form and function. Within our dataset, fishes show the weakest relationship between mandibular properties and overall feeding mode, with different feeding modes overlapping extensively in both jaw shape (Figure 5C) and function (Figure 2C). In contrast, feeding modes in salamanders and turtles, although still messy, show clearer trends in jaw speed *versus* strength (Figures 2D–E, 5D–E). When considering the architecture of the overall feeding mechanism, fishes stand out among vertebrates for the many kinetic elements typically involved in this system, including the cranium, jaws, hyobranchial apparatus, suspensorium, operculum, and pectoral girdle (*1, 2, 4–8, 11, 17, 18, 27, 39, 68, 74, 82*). Conversely, feeding-related movement beyond the jaws and hyobranchium is comparatively minimal to absent among salamanders and especially turtles (*10, 19, 27, 39, 74*), such that the overall feeding mechanism has fewer ‘degrees of freedom’ compared to fishes (*74*). The mandible in these latter clades is therefore both more directly tied to feeding performance (as per *65*) and more strongly subject to (i.e., less able to evade) performance trade-offs (as per *74*), hence a stronger coupling between mandibular properties and overall feeding mode. Supporting this interpretation, dietary modes in mammals—a clade in which the lower jaw is the only kinetic component of the skull (*68*)—show a particularly strong relationship with mandibular shape and mechanical advantage (see e.g., *47, 48, 83*).

Beyond these varied patterns of morphofunctional decoupling, complexity also arises in the methods typically used to quantify feeding performance. Although functional metrics (e.g., *2, 3–5, 30–32, 34, 36, 44, 45, 50, 51*, and the present study) and digital modelling (e.g., *2, 4, 33, 34, 36, 38, 44, 50, 51*) indicate what an animal is *capable* of performing, its *actual* behaviour does not necessarily realize this full range (*6, 21, 30, 40, 84*). Compounding this caveat, functional metrics likely provide a deceptively simple snapshot of actual feeding performance. For example, mechanical advantage is nearly ubiquitous among quantitative jaw studies (see e.g., *2, 3–5, 14, 31, 32, 37, 39, 40, 44, 45, 48, 50, 71, 85*), reflecting widespread acceptance of jaws as a lever system constrained to a simple speed-*versus*-force-transmission trade-off (*2, 4–6, 30, 36, 37, 39–43, 45, 48–51, 71, 73*). However, studies testing this assumption have found a more complex signal, with mechanical advantage showing inconsistent correlation with bite force in piranhas and pacus (*85*) and no relationship with the speed of mandibular depression in serranid fishes (*86*). Furthermore, this theoretical speed/strength trade-off assumes a quasi-static lever system, whereas in reality many biological structures comprise dynamic systems in which this trade-off has been mathematically invalidated (*87*). As another example, rotational efficiency was recently developed as a novel metric for jaw speed (*36*), but within our dataset often provides a very different signal from other speed-related metrics (Figures 2–3 and 8), suggesting further complexity in how these presumed proxies actually relate to ‘true’ jaw velocity.

*In vivo* studies can help to address these methodological shortcomings, as kinematic patterns do tend to differ consistently and predictably among different feeding modes (*6, 7, 11, 18, 19, 24, 26, 27, 88*). However, such studies are not a panacea for deciphering feeding modes: fishes often modulate their feeding behaviours based on flow properties (*70*), sensory input (*75*), type of prey (*84, 89*), and satiation (*90*), such that a single trial or study may not capture an animal’s full feeding repertoire (see also *1, 69, 91*); and, kinematic studies are inherently limited to living organisms, thus excluding the fossils and museum specimens critical to large-scale macroevolutionary analyses. In such cases, exploring other functional metrics (see e.g., *4, 36*) or other anatomical structures (e.g., the hyoid apparatus; *7, 12, 26, 28*) can help to extend our frame of reference beyond the mandible.

Despite these caveats to the typical utilization and conceptualization of mandibles in deciphering feeding modes, the lower jaw nonetheless remains a key means of insight into the evolution of vertebrate feeding. For example, although our finding of different morphofunctional trends in extant fishes, salamanders, and turtles is problematic when trying to apply a single biomechanical framework universally across vertebrates, this result is equally compelling in highlighting taxon-specific solutions to the challenge of accomplishing different feeding modes, thus emphasizing a need for more detailed investigation into form/function relationships, modularity, and decoupling within each group. Our finding of different morphofunctional trends in extant *versus* extinct taxa also highlights the continued importance of the fossil record in understanding macroevolutionary dynamics. Whereas extant vertebrate diversity is often envisioned as the result of progressive winnowing of an ancestrally much broader range of organismal form and function (e.g., ‘early burst’ models of disparity during adaptive radiations; *35*), our data instead suggest that early-evolving taxa within several vertebrate clades are in fact relatively restricted in both morphospace (Figure 5A, Figure S1) and mechanospace (Figure 2A, Figure S2), with crown-group representatives expanding into more varied morphologies and functions, thus expanding upon signals previously recognized among fishes (*35, 51*). The often-non-overlapping position of extinct *versus* extant taxa in morphospace (Figure 5A,C–E) and mechanospace (Figure 2A,C–E) further illustrates the intrinsic value of fossils in exploring combinations of form and function not seen in modern taxa, thus expanding our biomechanical and macroevolutionary frame-of-reference throughout deep time (see also *37, 92*). We emphasize, though, that accurate interpretation of these trends requires careful morphofunctional contextualization: as for any complex anatomical system, the mandible ultimately constitutes just one module within the broader network of bones, soft tissues, and behaviour defining a given feeding mechanism, which, although necessary to understanding the evolution of this overall system, is not innately sufficient to recapitulate such trends in their entirety.

### 3.2. Implications for early tetrapod evolution

In a study using many of the same functional metrics and early tetrapod taxa analyzed herein, Anderson, Friedman and Ruta (*32*) found amniotes to be the only group exploring novel mandibular functions, a signal which they interpreted as evidence for feeding modes remaining static until the advent of herbivory, quite late in the water-to-land transition (see also *44, 45*). However, our data impel reinterpreting these findings. Our analyses of mandibular form and function demonstrate that, although these lines of evidence can *sometimes* capture major transitions in overall feeding mode (e.g., the evolution of specialized biting in triggerfishes and parrotfishes), in most cases different feeding modes are indistinguishable at a vertebrate-wide scale (Figures 2 and 5). Therefore, although the changes in mandibular function and disparity previously recovered in amniotes *do* conceivably reflect their transition to herbivory, the *lack* of functional transformation in other taxa does *not* necessarily reflect overall stasis in feeding mode.

That the most extensive exploration of mechanospace (Figure 2) and morphospace (Figure 5) occurs among certain specialized biters has both biological and methodological bases. Beyond suggesting differing levels of morphofunctional constraint across different feeding modes—and, specifically, a release of constraint on biters (see also *7, 44, 45, 82*)—this greater diversification of mandibular form and function in dedicated biters also reflects how heavily these taxa rely upon the mandible (compared to e.g., suction-feeders, in which the hyobranchium plays an equal if not larger role in driving feeding performance; see e.g., *9, 12, 20*). This more dedicated reliance upon the mandible in turn presents an evolutionary impetus promoting diversification of this structure toward a more ‘optimal’ condition (see also 49, and e.g., echoing trends recognized in molluscivorous carnivorans by *83*), in this case one maximizing strength (Figure 2) and robustness (Figure 5). In contrast, the lack of mandibular diversification among other taxa *could* reflect a lack of diversification in feeding mode, but could just as reasonably reflect the one-to-many mapping pervasive throughout our dataset, in which similar mandibular shapes and/or functional traits occur in feeding mechanisms that are otherwise strikingly divergent.

The position of early herbivores within our morpho- (Figure 5) and mechanospaces (Figure 2) illustrates this latter concept. In mechanospace, diadectids and *Edaphosaurus* are positioned close to turtles—especially aquatic and semi-aquatic taxa such as sea turtles, softshells, and various testudinoids—reflecting a general emphasis on symphyseal length, adductor fossa length, and mechanical advantage (Figure 2A). However, although many of these extant taxa share with early herbivores a reliance upon jaw prehension (*93–96*), aquatically-feeding turtles also routinely exhibit both compensatory and inertial suction (*22, 93–95, 97*), with softshells often being entirely reliant upon the latter (*28, 98*). Notably, though, this capacity for suction generation is driven largely by the hyobranchial apparatus (*9*), while herbivory relies heavily on tooth morphology (*72*), such that analyses based entirely upon mandibles fail to capture these critical differences in overall feeding mode. Reinforcing this one-to-many mapping, this region of mechanospace also contains various suction-feeding and biting fishes, again illustrating that similarity in mandibular function does not necessarily indicate similarity in overall feeding mode, and should therefore not be interpreted as evidence of the *overall feeding mode* ‘remaining static’ across the origin of tetrapods.

Although mandibles cannot reliably predict transformations in *overall* feeding mode, jaw form and function *can* nonetheless provide key insight into how this structure *as a specific component of the early tetrapod body plan* evolved across the water-to-land transition. Rather than reflecting a simple shift from ‘fast’ to ‘strong’ jaws—traits typically assumed to characterize suction-feeding *versus* biting (see e.g., *1, 23, 36, 37, 44, 51, 60, 71, 72*), itself a transition widely portrayed as occurring across the origin of tetrapods (see e.g., *12, 13, 16, 33, 44, 45, 59–63*)—our data instead indicate a more complex evolutionary trajectory. Whereas taxa on the outer bounds of this transition (i.e., early-diverging tetrapodomorph fishes such as osteolepidids, and later-diverging stem- and crown-amniotes) exhibit robust (Figure 5F, Figure S1F) and generally strong (Figure 2F, Figure S2F) jaws, a phase of highly gracile mandibles (Figure 5F) with little maximization of any functional traits (Figure 2F) occurs among late-diverging tetrapodomorph fishes, early tetrapods, and early-diverging amniotes and temnospondyls. This trend evokes that recovered by Dickson, Clack, Smithson and Pierce (*57*) for the evolution of the tetrapod humerus, whereby early tetrapods occupy a ‘performance valley’ reflecting a compromise in locomotor efficiency between swimming in water and walking on land. Our mandibular data point to a similar ‘valley’ of jaw performance across the origin of tetrapods (*contra* hypotheses of stasis in e.g., *32, 44*). To what extent these changes would have impacted the broader feeding mechanism—i.e., whether overall feeding performance was concomitantly ‘sub-optimal’, or was instead mitigated by transformations elsewhere to the feeding apparatus—ultimately requires quantifying trends in other feeding-related structures, such as the hyoid apparatus and cranium. Intriguingly, the lack of clear differentiation in the mandibular requirements for different feeding modes hints at a complex performance landscape across this transition, with the multiple biomechanical factors and evolutionary pressures associated with feeding enabling multiple adaptive peaks (see also *43, 49*). Ultimately, the ability of similar mandibular shapes (Figure 5) and functions (Figure 2) to be leveraged across diverse environments and feeding strategies likely facilitated evolution across this landscape, adding a new dimension to our understanding of the complex evolutionary dynamics underlying this transition.

Finally, the different manifestations of jaw shape (Figure 5) and function (Figure 2) in extant *versus* extinct taxa present a salient challenge to deciphering form and function throughout the fossil record. Fossils are often interpreted through the lens of modern biota, such as via Extant Phylogenetic Bracketing (*99*). However, these approaches depend upon extant taxa constituting a valid empirical framework. Our data instead show that key trends manifest differently across major vertebrate partitions, and that fossils—especially those representing particularly deep divergences—often fall entirely outside modern patterns. Our linear discriminant analyses exemplify this finding (Figures 3–4): beyond having generally low accuracy (Table 1), these analyses struggle to make reasonable predictions for fossils (Tables S4–5), consistently misclassifying taxa of widely-accepted feeding modes (e.g., the herbivorous diadectids and *Edaphosaurus* almost always being predicted as ‘hybrid-‘ or even ‘suction-feeders’) or omitting certain feeding modes altogether (i.e., models trained based on salamanders, a clade which lacks any ‘pure’ suction-feeders). Reconstructing feeding across the tetrapod water-to-land transition is especially problematic given that early tetrapods follow different morphofunctional trends than extant vertebrates (Figures 2 and 5), reinforcing that the mandible alone is insufficient for determining feeding modes in extinct taxa. Although lower jaws represent a critical *piece* of evidence in drawing higher-level interpretations of feeding mode, the integration of broader lines of evidence—such as other feeding-related structures (e.g., the hyobranchial apparatus; *7, 8, 12, 26, 28*) or analytical approaches (e.g., network analysis, as by *62,* or computational fluid dynamics, as by *78*)—will be necessary to draw rigorous and accurate conclusions.

### 3.3. A new paradigm?

The nuances illuminated by our data bolster recent calls (see e.g., *54*) to move beyond the “suction-feeding *versus* biting” and “speed *versus* strength” paradigm that has dominated research into the evolution of vertebrate feeding, from the origin of tetrapods (e.g., *44, 59*) to the advent of jaws themselves (e.g., *31, 36*) to large-scale trends in vertebrate feeding ecology (e.g., *5*). In the context of vertebrate water-to-land transitions, logically any evolutionary shift between these habitats must, at some level, entail a transition from suction generation to jaw prehension for prey capture, given the widespread usage of suction-feeding in aquatic environs but its fundamental ineffectiveness on land (*8, 9, 12, 18, 44, 45, 54, 59*). However, reducing feeding ecology to this nearly-dichotomous perspective almost certainly elides the complexity involved in such transitions. The question at play here is ultimately twofold: does a strict trade-off between these overall feeding modes exist in the first place and, if so, is this trade-off applicable to events such as the origin of tetrapod terrestriality?

Under a true trade-off, suction-feeding and biting would constitute two discrete categories bearing opposing functional demands (see e.g., *36, 44, 51*), with taxa being specialized for either one mode or the other (see e.g., *59*) and a transition across this dichotomy requiring an inherent decrease in performance as certain functional traits are sacrificed in favour of others (*1, 43, 67, 74, 100, 101*). However, our data highlight that this functional conflict does not necessarily exist—at least when considering the mandible—as these feeding modes instead overlap in both jaw shape (Figure 5) and function (Figure 2). Indeed, as has been well recognized throughout both modern taxa (e.g., *1, 2, 3, 6, 12, 17, 23*) and fossils (e.g., *54, 60, 61*), ‘hybrid’ feeding modes incorporating both suction feeding and jaw prehension are common throughout vertebrates. Although this combined usage *may* entail a decrease in performance (e.g., a reduction in suction efficacy in some post-metamorphic salamanders; *102, 103, 104*) (see also *8, 24, 26*), taxa such as teleosts (*88, 100*), gars (*17, 54*), lungfishes (*20, 105, 106*), sirenids (*107*), and newts (*8, 108*) instead demonstrate that these strategies can coexist without either necessarily needing to be ‘sacrificed’. For example, suction performance can remain consistent regardless of reliance on biting among teleost fishes, reflecting a lack of trade-off between these feeding modes (*88, 100*). In the context of early tetrapod evolution, the evolutionary advantages afforded by this versatility (see also *49*) were in fact likely critical to traversing the water-to-land boundary.

In light of these myriad ‘intermediate’ feeding modes, a more accurate conceptualization of this transition might be to consider it as a continuum, rather than a dichotomy (as suggested by e.g., *2, 54, 61*, and implied by the Pareto approach taken by *36, 44, 51*). However, even this framework is not entirely satisfying given the tremendous morphofunctional variation encapsulated at the supposed endpoints of this transition. For example, although ‘biting’ may seem straightforward, simply reflecting reliance on jaw prehension (*1*), this strategy in reality takes numerous forms, including slicing (e.g., piranhas (*23, 72, 85*)), scraping and/or excavating (e.g., parrotfishes (*23, 42, 109*)), crushing and/or chewing (e.g., lungfishes (*20, 105, 106*); pacus (*72, 81, 85*); salamanders (*10, 12, 107*)), lateral snapping (e.g., gars (*17*)), scale-eating (e.g., cichlid and serrasalmid fishes (*23, 79, 85*)), pharyngeal biting (e.g., moray eels (*110*); cichlids (*71*)), and combined jaw and lingual prehension (e.g., many terrestrial turtles (*9, 98, 111*); and terrestrial salamanders (*12, 24*)). Each instantiation of ‘biting’ could conceivably have its own manifestation of optimal form and function—for example, carnivorous piranhas have a greater bite force, larger adductor muscles, different mechanical advantage, and lower articular offset than the closely related, herbivorous pacus, despite both groups being ‘biters’ (*85*) (see also *23, 42, 79, 109*)—each of which could in turn ‘trade off’ with suction-feeding in a distinctive manner. The transition between these feeding modes therefore likely entails countless morphofunctional paths, each with its own macroevolutionary dynamics, reflecting a complex and multi-peaked adaptive landscape (see also discussion in *54*) rather than a single ‘axis’ of evolution between two fixed endpoints.

The applicability of a ‘suction-feeding *versus* biting’ trade-off to the origin of tetrapods is further undermined by the tenuous connection between these feeding modes and a given organism’s ecology. For instance, whereas biting is clearly well-suited to feeding on land, many aquatic taxa—from piranhas and parrotfishes (*23, 85*), to sirenid salamanders (*10, 12, 107*), to sea turtles (*73, 96*), among others (see Table S1)—also rely upon jaw prehension when feeding in water. Therefore, even if a transition from suction-feeding to biting *did* occur in early tetrapods, this does *not* necessarily correspond to the transition to land. Instead, adaptations for biting-based feeding could very well have arisen in taxa that were still fundamentally aquatic, either preceding (*60*), coinciding with (e.g., *Tiktaalik*, *Acanthostega*; *16, 54, 59*), or shortly following (e.g., *Whatcheeria*, *Crassigyrinus*; *33, 61, 64*) the origin of tetrapods (see also *8*). Furthermore, various *other* terrestrial feeding mechanisms exist in addition to—or, in some cases, instead of—biting, such as lingual feeding in turtles (*9, 98, 111*) and salamanders (*24*); indeed, certain early tetrapods have been hypothesized to have used feeding modes beyond the typical continuum of ‘aquatic suction-feeding’ to ‘terrestrial biting’, such as filter-feeding (*Spathicephalus*; *61, 112*), aquatic durophagy (*Acherontiscus*; *61, 113*), or complex chewing (*12*). Given this ecofunctional complexity, conceptualizing the origin of tetrapods exclusively within the framework of ‘suction-feeding *versus* biting’ therefore risks ignoring the possibility of diversification into other feeding mechanisms across this transition.

Thus, although the notion of a transition from suction-feeding to biting across the origin of tetrapods (and other vertebrate ecomorphological transitions more broadly) may be useful as a guiding framework, it should ultimately be taken as just that: a guide, with the biological reality of this transition involving much more nuance and uncertainty. Rather than being limited to a preconceived framework of specific feeding mechanisms or trade-offs, focusing instead on larger trends in morphology (e.g., evolutionary rates; *37, 114*), function (e.g., functional metrics; *14, 37, 39*), and morphofunctional diversification (e.g., patterns of disparity; *31, 32, 35, 37, 39*) will permit a more agnostic perspective when quantifying major transformations in vertebrate feeding.

## 4. Methods

### 4.1. Taxon Sample

#### 4.1.1. Taxon selection

To assess vertebrate-wide trends in mandibular form and function as related to suction-feeding *versus* biting, we compiled an ecomorphologically and phylogenetically diverse dataset of lower jaws from 214 species, comprising 95 extant and 119 extinct taxa (Figure 1A; Table S1). Our modern sample contained 34 fishes, 22 salamanders, and 39 turtles, representing a range of feeding modes (see below). Because these extant taxa served as a known ‘baseline’ against which to validate long-held assumptions, we therefore focused on species with well-characterized feeding modes, especially those having undergone detailed biomechanical investigation or having been used in other studies examining transitions in feeding mode (e.g., *Megalops* and *Micropterus* as common exemplars of suction-feeding; see e.g., *6, 7, 11*). Our extinct sample focused on species spanning the tetrapod water-to-land transition, from tetrapodomorph fishes (n = 14) through early tetrapods (n = 14) to temnospondyls (n = 6) and early amniotes (n = 19 stem, 5 crown). We also sampled extinct members of the various extant clades in our dataset—i.e., stem-Osteichthyes (n = 1; grouped with stem-sarcopterygians following *37*), actinopterygians (n = 10 stem, 14 crown), sarcopterygian fishes (n = 3 stem, 22 crown), salamanders (n = 1 stem, 1 crown), and turtles (n = 4 stem, 5 crown)—to provide evolutionary context for these groups. To assess trends both within and across major vertebrate groups, we performed each analysis (see below) for fourteen different sub-partitions within our overall dataset: all taxa, extant taxa, extinct taxa, all fishes, extant fishes, extinct fishes, all salamanders, extant salamanders, all salamanders and temnospondyls, extinct salamanders and temnospondyls, all turtles, extant turtles, extinct turtles, and extinct taxa spanning the water-to-land transition (i.e., tetrapodomorph fishes, early tetrapods, temnospondyls, and stem- and crown-amniotes). For simplicity, we focused our Results and Discussion on only the most informative analyses, but provide data for all taxon partitions in the Supplementary Materials.

#### 4.1.2. Feeding mode classification

Based on published descriptions of feeding behaviour and/or biomechanics (Table S1), we classified each extant species as either a ‘suction-feeder’ (i.e., relying entirely upon suction to capture and ingest prey; n = 16), a ‘biter’ (i.e., relying entirely upon jaw prehension during prey capture; n = 28), or a ‘hybrid-feeder’ (i.e., routinely using both suction and biting, either concurrently during a single feeding event or separately during distinct feeding events; n = 51). We also assigned feeding modes to extinct taxa where possible (Table S1). To keep these categories rigorous and internally consistent, we only assigned feeding modes to fossils for which previous authors had clearly addressed the role of both suction-feeding and biting. For example, the Mesozoic holostean *Scheenstia* has been described as durophagous (*80*), indicating a reliance upon biting; however, extant durophages also commonly employ suction (e.g., lungfishes; *11, 20, 105, 106*), such that this conclusion of ‘durophagy’ alone is not sufficient to distinguish between a designation as a ‘biter’ or a ‘hybrid-feeder’, hence our classification of this taxon as ‘unknown’. Ultimately, we were able to assign feeding modes to 65 of 119 fossils (n = 9 suction-feeders, 33 hybrid-feeders, 23 biters, 54 unknown).

Recognizing that any categorization of feeding modes inevitably requires arbitrary sub-division of an inherently continuous spectrum of behaviours, we also explored the impact of using a more detailed classification scheme. Specifically, we additionally scored each taxon on a scale of 1–5, where ‘1’ and ‘5’ correspond to the above definitions of suction-feeding and biting, respectively, and ‘2’–‘4’ reflect more precise definitions of ‘hybrid-feeding’: ‘2’ reflects an emphasis on suction generation with a minor role of biting (n = 10 extant, 5 extinct), ‘4’ represents an emphasis on biting with a minor role of suction-feeding (n = 22 extant, 14 extinct), and ‘3’ represents equal reliance on both feeding modes (n = 19 extant, 14 extinct) (see Table S1 for classifications). However, because these alternative classifications had little effect on the key trends recovered in our analyses, we focus herein on the simpler, 3-mode scheme, with results from the 5-mode classification being presented in the Supplementary Materials (see Figures S3–8).

#### 4.1.3. Phylogeny

To provide evolutionary context for our data, we compiled a ‘super-tree’ of the taxa in our dataset (Figure 1A; Data S1) using *TimeTree* [v.5] (*115*) for extant taxa and various published phylogenies for fossils (Data S2). We assembled and time-calibrated the tree in R [v.4.3.2] (*116*) and RStudio [v.2023.12.1] (*117*), with the latter performed using the function “*timePaleoPhy*” in the R package *paleotree* [v.3.4.5] (*118*) and with first-appearance and last-appearance dates for extinct taxa obtained from the Paleobiology Database (*119*).

### 4.2. Data Collection

#### 4.2.1. Silhouette generation

For all extant and some extinct species, we generated silhouettes from 3D models of the mandible (Table S1). We used the left hemimandible for all specimens, unless it was damaged, in which case we used a mirrored version of the right hemimandible. For taxa lacking separate hemimandibles (e.g., turtles), we digitally split the jaw vertically along the mandibular symphysis. Data were sourced from online repositories (MorphoSource; AdMorph), individual publications, or micro-computed tomography (microCT) or photogrammetric scans (see Table S1 for full information regarding specimen sources). Some specimens were already available as mesh files; in other cases, we generated the meshes ourselves, either from microCT data which we segmented in Mimics [v.24.0–v.27.0] (*120*), or via photogrammetry using a variety of software (RealityScan [v.2.0.1] (*121*); MeshLab [v.2022.02] (*122, 123*); Agisoft PhotoScan Standard [v.1.4.1.5925] (*124*); CloudCompare [v.2.13.2] (*125*)).

We post-processed each mesh in 3-matic [v.16.0–v.20.0] (*126*). For each specimen, we generated two final meshes: a ‘raw’ hemimandible, which we aligned to a consistent reference position (aligned along its inertial axes, with +X pointing medially, +Y pointing dorsally, and +Z pointing posteriorly) but otherwise left unaltered; and a ‘processed’ hemimandible, which we modified by removing teeth (where applicable), repairing any mesh defects, smoothing and simplifying the overall mesh, and aligning it into the aforementioned reference position. We used both the raw and processed meshes to collect functional data, and the processed mesh to collect morphological data. To generate the silhouette needed for the latter, we imported a lateral screenshot of each processed mesh into Adobe Photoshop and converted it into a black-and-white silhouette, which we exported in JPG format (Data S3).

For the remaining taxa in our dataset, for which 3D data were unavailable, we instead generated silhouettes from published figures (see Table S1 for image sources). As for the 3D-based silhouettes, we used the left hemimandible where possible, or a mirrored right hemimandible when the left was not sufficiently depicted in the associated publication. We used Adobe Photoshop to post-process each published figure (specifically, to remove teeth where applicable) and to convert it into a black-and-white silhouette, which we exported in JPG format (Data S3).

#### 4.2.2. Geometric morphometrics

We quantified the shape of each silhouette via elliptical Fourier analysis (EFA), as commonly used in recent studies of jaw evolution (e.g., *34, 35, 36, 44, 45, 50, 51*) and as implemented in the R package *Momocs* [v.1.4.1] (*127*).

Prior to performing EFA, we first used *tpsDig* [v.2.31] (*128, 129*) and *tpsUtil* [v.1.83] (*128, 130*) to place a set of fixed landmarks on each silhouette. These landmarks were not themselves used for geometric morphometric analysis, but instead served to ensure consistent orientation of each silhouette during Procrustes superimposition. We placed four landmarks, at: [1] the anterodorsal tip of the mandibular symphysis; [2] the dorsalmost point of the coronoid process (placed at the midpoint of the process if it lacked a distinct dorsalmost point, or at the position between the tooth row and jaw joint where the mandible was deepest if lacking a distinct coronoid process altogether); [3] the centre of the jaw joint; and [4] the posterior tip of the retroarticular process. After saving the landmark files in TXT format, we then imported them into R alongside the corresponding silhouettes, using a modified version of the *Momocs* function “*tie_jpg_txt*”.

Following Tseng, Garcia-Lara, Flynn, Holmes, Rowe and Dickson (*50*), we next used various functions in *Momocs* to perform a series of adjustments on the silhouettes, thus ensuring comparable formatting of each image prior to EFA: after smoothing each silhouette for ten iterations (using the function “*coo_smooth*”), we centred all silhouettes on the origin (“*coo_center*”) and sub-sampled each silhouette to 500 outline points (“*coo_sample*”). We then subdivided our overall silhouette dataset into the various taxon partitions noted above and performed generalized Procrustes analysis for each partition using the function “*fgProcrustes*”, with the fixed landmarks from *tpsDig* used to guide this alignment. Finally, we performed EFA on each taxon partition using the function “*efourier*”, with a harmonic power of 99% (i.e., using the number of harmonics required to capture 99% of the shape variation in the silhouettes).

#### 4.2.3. Functional metrics

To assess trends in jaw function, we measured eight metrics commonly used in previous studies of mandibular evolution (Figure 1B), which together represent proxies for jaw strength (mechanical advantage, relative adductor fossa length, relative symphyseal length), jaw speed (maximum inverse aspect ratio, average inverse aspect ratio, rotational efficiency), or other aspects of jaw function (relative articular offset, symphyseal angle).

**Mechanical advantage** (“MA”) reflects the efficiency of bite force transmission, measured as the ratio of the jaw-closing in-lever (i.e., the distance from the centre of the jaw joint to an estimated position of adductor muscle insertion) divided by its out-lever (measured as the distance from the centre of the jaw joint to the anterior tip of the jaw) (see e.g., *2, 3–5, 14, 31, 32, 37, 39, 40, 44, 45, 48, 50, 85*). Following various studies (e.g., *2, 3, 39, 41, 50*), we generally treated the dorsal-most tip of the coronoid process as the location of adductor muscle insertion, and thus the anterior endpoint of the MA in-lever. When the coronoid process was approximately level, we instead measured to its dorsal midpoint. When a specimen lacked a distinct coronoid process (e.g., as in many extinct taxa), we measured to the position between the tooth row and jaw joint where the jaw was vertically deepest; following Anderson (*14*) and Anderson, Friedman, Brazeau and Rayfield (*31*), this position approximates where the second moment of area is highest, thus where the jaw is stiffest, thus where the adductor musculature is most likely to principally act.

**Relative adductor fossa length** (“AF”) acts as a proxy for the cross-sectional area of the adductor musculature, which is in turn proportional to its force output (see e.g., *31, 32, 45*). We calculated this metric as the length of the adductor insertion region divided by the overall horizontal length of the jaw. In most taxa, the adductor fossa is an obvious opening on the dorsal surface of the mandible, in which case we measured ‘adductor fossa length’ as the distance from the centre of the jaw joint to the anterior terminus of the adductor fossa. However, in some taxa, the adductor insertion region is shifted onto either the medial (as in many fishes) or lateral (as in turtles) surface of the mandible; in such cases, we measured ‘adductor fossa length’ as the distance from the jaw joint to the dorsal ‘apex’ of the adductor insertion region. In other taxa, a distinct adductor fossa or insertion region was not readily visible; in such cases, we measured ‘adductor fossa length’ as the distance from the jaw joint to either the anterior terminus of the coronoid process (reflecting the maximum potential region of adductor muscle insertion, similar to the approach taken by *31*) or the posterior terminus of the tooth row (in taxa such as early tetrapods which lack a distinct coronoid process; similar to the approach taken by *32*).

**Relative symphyseal length** (“SL”) is a proxy for symphyseal robustness and relates to the forces (e.g., shear stress, torsion) experienced during jaw use (see e.g., *31, 32, 45*) (see also *38*). We calculated this metric as the maximum length of the mandibular symphysis divided by the overall horizontal length of the jaw, with maximum symphyseal length being measured from the jaw tip to the posteroventral tip of the mandibular symphysis. In instances where the mandibular symphysis was not directly visible (e.g., silhouettes derived from lateral-only views of the jaw), we approximated the posteroventral tip of the symphysis as the ‘inflection point’ of the anterior terminus of the jaw, where the ventral margin transitions from descending roughly vertically to roughly posteriorly, following previous depictions of this metric (*31, 32*).

The **orientation or angle** of the mandibular symphysis (“SA”) is related to the forces (e.g., shear stress, torsion) experienced during jaw use (see e.g., *31*). We calculated this metric as the angle of the mandibular symphysis (measured along its maximum length) relative to the long axis of the jaw.

Aspect ratio (AR)—i.e., jaw height divided by jaw length—is a proxy for second moment of area, and thus flexural stiffness (*31, 32, 45*). Conversely, inverse AR—i.e., jaw length divided by jaw height—can instead act as a proxy for jaw velocity, based on the principle that more elongate mandibles can be displaced more readily than stouter ones (*50*). We calculated aspect ratio in two ways. First, we calculated **maximum inverse aspect ratio** (“maximum AR” or “mAR”), comprising the overall horizontal length of the mandible divided by its maximum vertical height (see e.g., *31, 32, 45, 50*). Because the maximum height of the mandible can be skewed by features such as a particularly tall coronoid process (as in e.g., *Polypterus*), we also calculated **average inverse aspect ratio** (“average AR” or “aAR”) (see e.g., *31, 32, 45*). We measured this metric as the overall horizontal length of the mandible divided by its average height, with the latter calculated as the silhouette’s surface area divided by its overall horizontal length.

**Rotational efficiency** (“RE”) is a recently developed metric reflecting the speed of jaw rotation given one unit of energy input (see e.g., *34, 36, 44, 51*). We calculated this metric as 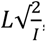, where *v* represents the velocity of the jaw tip, *L* represents the distance from the jaw tip to the centre of the jaw joint, and *I* represents the rotational inertia of the jaw. Rotational inertia was itself calculated as *mr*^2^, where *m* represents a proxy for jaw mass (in this case, the total surface area of the mandible) and *r* represents the distance from the mandible’s centre-of-mass to its centre-of-rotation (i.e., the centre of the jaw joint).

**Relative articular offset** (“AO”) reflects the type of tooth row occlusion: ‘scissor-like’ occlusion when the jaw joint is directly in line with the tooth row (typical of carnivores or other taxa with ‘slicing’ jaw movements) *versus* ‘vice-like’ occlusion when the jaw joint is offset from the tooth row (typical of herbivores and crushers) (see e.g., *14, 31, 32, 45, 47, 85*). We calculated this metric as the perpendicular distance between the centre of the jaw joint and a line running tangent to the pre-coronoid region of the jaw, divided by the maximum vertical height of the jaw. For most taxa, we represented the ‘tangent line’ using a line passing through the jaw tip and the posterior terminus of the tooth row. For taxa lacking teeth (e.g., turtles), we replaced the ‘posterior terminus of the tooth row’ with either the posterior terminus of the triturating surface or the anterior base of the coronoid process, such that a line connecting this point to the jaw tip would run roughly tangent to the pre-coronoid region of the jaw.

Finally, we also measured **normalized jaw length** (see e.g., *30, 31, 47*), reflecting overall horizontal length divided by the square root of the silhouette’s surface area. However, we ultimately excluded this metric from our analyses as it was extremely strongly correlated with average inverse aspect ratio (Figure S9).

#### 4.2.4. Collection of functional metrics

We designed a custom extension in Rhinoceros 3D [v.7.SR38] (*131*) to automate the collection of functional metrics, using the bundled visual programming environment Grasshopper [v.1.0.0007] (*132*) (Data S4). For mesh-based specimens, the Grasshopper macro first automatically converted each 3D mesh into a 2D object by projecting it onto its sagittal plane. The functional metrics and underlying measurements were then calculated using a combination of automatically-detected measurements (i.e., jaw length, height, and surface area) and manually-placed reference points (see Data S4 for details).

We designed a similar macro for the image-based specimens (Data S4). Because Rhino3D cannot directly import image files, we first converted each silhouette into a vector-format object using Adobe Illustrator (see Data S4 for details). Upon importing this Illustrator file into Rhino3D, we next used the Grasshopper macro to scale each silhouette to its true size and re-orient it into a reference position consistent with that used for mesh-based specimens (i.e., aligned along its inertial axes, with +Y pointing dorsally and +Z pointing posteriorly). As for the mesh-based specimens, functional metrics were then calculated using a combination of automatically-detected measurements and manually-placed reference points (see Data S4 for details).

After collecting these metrics in Rhino3D, we performed final adjustments and clean-up in R. Specifically, we used R to calculate normalized jaw length, convert aspect ratio into inverse aspect ratio, and convert RE into units of metres per second (Data S5).

### 4.3. Data Analysis

We performed all analyses in R [v.4.3.2] (*116*) and RStudio [v.2023.12.1] (*117*), using the scripts provided in Data S5.

#### 4.3.1. Correlation among functional traits

To test for trade-offs between strength- and speed-related traits, we used the function “*cor*” from the base-R package *stats* [v.4.3.2] (*116*) to calculate the Pearson correlation among the functional metrics within each taxon partition. We visualized the resulting correlation matrices as heatmaps using functions from the package *ggplot2* [v.4.0.2] (*133*), with statistical significance calculated using the function “*ggcorrmat*” from the package *ggstatsplot* [v.0.13.5] (*134*).

#### 4.3.2. Principal components analysis

To visualize the morphospace and mechanical space (hereafter, “mechanospace”) for each taxon partition, we performed principal component analyses (PCA) on the EFA outputs and functional metrics, respectively. For each PC plot, we colour-coded taxa according to both feeding mode and higher taxon, using functions from *ggplot2* [v.4.0.2] (*133*) and *ggConvexHull* [v.0.1.0] (*135*). We assessed the statistical significance of each grouping method using permutational multivariate analysis of variance (PERMANOVA), conducted via the function “*adonis2*” from the R package *vegan* [v.2.6-6] (*136*), using a Euclidean distance matrix and 10,000 permutations. For groupings with more than two categories, we also conducted a post-hoc pairwise PERMANOVA via the function “*pairwise.adonis2*” from the R package *pairwiseAdonis* [v.0.4.1] (*137*), again using a Euclidean distance matrix and 10,000 permutations.

#### 4.3.3. Cluster analysis

To assess which taxa were most similar in morphospace, we performed spectral clustering analyses on the PCA coordinates for each taxon partition, using the function “*Spectrum*” from the R package *Spectrum* [v.1.1] (*138, 139,* see also implementation by *140*). This algorithm assigns data-points into clusters using information about shared nearest-neighbours derived from a graph of the underlying data, with the optimal number of clusters determined based on the distribution of eigenvectors from this graph (*138*). Because all inputs to this spectral clustering algorithm are considered equally, PC axes that capture only small amounts of variance in the data become effectively ‘overweighted’ relative to more explanatory axes; as such, when running this cluster analysis, we used only PC axes with variance >5%. We visualized the resulting cluster assignments using functions from *ggplot2* [v.4.0.2] (*133*) and *ggConvexHull* [v.0.1.0] (*135*).

#### 4.3.4. Discriminant analysis

To assess trends in our functional data, we subjected our functional metrics to linear discriminant analysis (LDA), using functions from the R packages *klaR* [v.1.7-3] (*141*) and *candisc* [v.0.9.0] (*142, 143*). LDA recognizes pre-assigned groups within the dataset, so as to maximize the separation among these groups in the resultant multivariate space. Our goal in performing LDA was to test whether different feeding modes could effectively be ‘forced’ into different regions of mechanospace, and in turn whether the functional metrics underlying this distribution corresponded to assumed trade-offs in strength *versus* speed.

As is typical of discriminant analyses—and machine learning algorithms in general—LDA requires both a training dataset (i.e., taxa of known feeding mode, from which the multivariate model is constructed) and a test dataset (i.e., taxa of unknown feeding mode, which are then assigned into categories based on the aforementioned model). To assess trade-offs within each major taxon partition represented in our dataset, we performed LDA using eight different training datasets, each comprising the species of known feeding mode within that partition: all taxa (n = 160), extant taxa (n = 95), all fishes (n = 66), extant fishes (n = 34), all salamanders (n = 23), extant salamanders (n = 22), all turtles (n = 42), and extant turtles (n = 39). Using the resultant statistical models, we then predicted the feeding mode of all remaining taxa within our overall dataset, using leave-one-out cross-validation to determine the accuracy of each model’s predictions. We visualized the resulting discriminant spaces using functions from *ggplot2* [v.4.0.2] (*133*) and *ggConvexHull* [v.0.1.0] (*135*).

#### 4.3.5. Allometry

To assess the influence of allometric scaling in our dataset, we examined the effect of size on both shape and function. To represent size, we used the log_10_-transformed square root of the surface area of each silhouette. (Although *Momocs* reports the centroid size for each silhouette, this value simply reflects the raw size of each inputted image and therefore is not biologically meaningful.) To represent shape and function, we used the scores from PC axes 1–3 from our morphospace and mechanospace analyses, respectively. Using these proxies, we performed both Ordinary Least Squares and Phylogenetic Generalized Least Squares regression. We performed the former via the function “*lm*” from the base-R package *stats* [v.4.3.2] (*116*) and the latter via the function “*gls*” from the R package *nlme* [v.3.1-164] (*144, 145*) alongside functions from the packages *ape* [v.5.8-1] (*146, 147*), with the results plotted using functions from *ggplot2* [v.4.0.2] (*133*).

### 4.4. Abbreviations

#### 4.4.1. Functional metrics

Functional metrics are labelled in Figures 1–3 and 8 using the following abbreviations: *aAR*, average inverse aspect ratio; *AF*, relative articular offset; *AO*, relative articular offset; *MA*, mechanical advantage; *mAR*, maximum inverse aspect ratio; *RE*, rotational efficiency; *SA*, symphyseal angle; *SL*, relative symphyseal length.

#### 4.4.2. Taxa

Taxa are labelled in Figures 2, 3, 5, and 6 using the following abbreviations: Aca, *Acanthostega*; Ach, *Achoania*; Ado, *Adocus*; Alb, *Albula*; Ami, *Amia*; Amp, *Amphiuma*; Amy, *Amyda*; And, *Andrias*; Ang, *Anguilla*; Ant, *Anthracosaurus*; Apa, *Apalone*; Arc, *Archeria*; Atr, *Atractosteus*; Bal, *Balistes*; Bir, *Birgeria*; Bol, *Bolitoglossa*; Bue, *Buettnererpeton*; Cac, *Carettochelys*; Cae, *Caerorhachis*; Cap, *Capitosaurus*; Car, *Caretta*; Cat, *Catoprion*; Cet, *Cetoscarus*; Chd, *Chelydra*; Che, *Chelonoidis*; Chi, *Chitra*; Chl, *Cheilinus*; Chp, *Chirodipterus*; Chr, *Chersina*; Cht, *Chitala*; Col, *Colosteus*; Cra, *Crassigyrinus*; Cry, *Cryptobranchus*; Cte, *Ctenolabrus*; Cuo, *Cuora*; Dap, *Dapedium*; Den, *Dendrerpeton*; Der, *Dermochelys*; Des, *Densignathus*; Dia, *Diadectes*; Dic, *Dicamptodon*; Dil, *Diplocercides*; Din, *Diandongichthys*; Dip, *Dipnorhynchus*; Dit, *Dipterus*; Dog, *Dogania*; Dor, *Doragnathus*; Dur, *Durialepis*; Ech, *Echidna*; Ect, *Ectosteorhachis*; Eda, *Edaphosaurus*; Eld, *Eldeceeon*; Elg, *Elginerpeton*; Eoh, *Eoherpeton*; Eot, *Eothyris*; Eso, *Esox*; Eus, *Eusthenopteron*; Gec, *Geoclemys*; Gep, *Geophagus*; Ges, *Gephyrostegus*; Gly, *Glyptops*; Gog, *Gogosardina*; Gon, *Gogonasus*; Gop, *Gopherus*; Gre, *Greererpeton*; Gui, *Guiyu*; Gym, *Gymnothorax*; Heo, *Heosemys*; Hol, *Holodipterus*; Hom, *Homopus*; Hop, *Holoptychius*; How, *Howqualepis*; Hyb, *Hynobius*; Hyd, *Hydrolycus*; Hyn, *Hyneria*; Ich, *Ichthyostega*; Jar, *Jarvikina*; Ken, *Kenichthys*; Ket, *Kentuckia*; Kuw, *Kuwavaatakdectes*; Lat, *Latimeria*; Led, *Lepidochelys*; Lem, *Lepomis*; Lep, *Lepidosiren*; Lim, *Limnoscelis*; Lis, *Lissemys*; Man, *Manouria*; Mar, *Marmorerpeton*; Mec, *Megalocephalus*; Med, *Medoevia*; Mee, *Meemannia*; Meg, *Megalops*; Mem, *Megamastax*; Met, *Metaxygnathus*; Mic, *Micropterus*; Mim, *Mimipiscis*; Mon, *Monacanthus*; Nec, *Necturus*; Neo, *Neoceratodus*; Nep, *Neopteroplax*; Neu, *Neusticemys*; Nga, *Ngamugawi*; Nic, *Nichollsemys*; Oro, *Orobates*; Oso, *Osorioichthys*; Ost, *Osteolepis*; Oxy, *Oxycheilinus*; Pac, *Pachytriton*; Pam, *Parmastega*; Pan, *Panderichthys*; Par, *Paramesotriton*; Pec, *Pelochelys*; Ped, *Pelodiscus*; Phl, *Pholiderpeton*; Pho, *Phonerpeton*; Pla, *Platysternon*; Ple, *Plethodon*; Pls, *Plastomenus*; Plt, *Platycephalichthys*; Pol, *Polypterus*; Por, *Portlandemys*; Pow, *Powichthys*; Pro, *Protocaptorhinus*; Prt, *Proterogyrinus*; Psa, *Psammobates*; Pse, *Pseudotriton*; Ptr, *Protorothyris*; Pyg, *Pygocentrus*; Pyx, *Pyxis*; Reg, *Regalerpeton*; Rhc, *Rhinochelys*; Rhi, *Rhinoclemmys*; Rom, *Romeria*; Sal, *Salamandrina*; Sau, *Saurichthys*; Sax, *Saxochelys*; Sca, *Scarus*; Sch, *Scheenstia*; Ser, *Serrivomer*; Sey, *Seymouria*; Sig, *Sigournea*; Sil, *Silvanerpeton*; Sim, *Simenchelys*; Sir, *Siren*; Spe, *Speonesydrion*; Spo, *Spodichthys*; Ste, *Sternotherus*; Stk, *Stenokranio*; Stl, *Styloichthys*; Sty, *Stylemys*; Syn, *Synaphobranchus*; Teg, *Tegeolepis*; Tes, *Testudo*; Thi, *Thiollierepycnodus*; Tik, *Tiktaalik*; Tri, *Trionyx*; Trm, *Trimerorhachis*; Tse, *Tseajaia*; Tun, *Tungsenia*; Tyl, *Tylototriton*; Ura, *Uranoscopus*; Url, *Uranolophus*; Ven, *Ventastega*; Wat, *Watsonulus*; Wes, *Westlothiana*; Wha, *Whatcheeria*; Xen, *Xenentodon*; Yel, *Yelangichthys*; Yme, *Ymeria*; You, *Youngolepis*.

## Acknowledgments

**Acknowledgments:** We thank C. Byrd, A. Kowalczyk, and S. Kennedy-Gold (MCZ) for their assistance with specimen access, R. Figueroa for advice on taxon sampling and access to microCT data, H. Nguyen for assistance with segmentation of microCT scans, P. Bishop for assistance with photogrammetry protocols, and various members of the Pierce Lab for helpful discussions during project development.

## Author contributions

***Conceptualization***: C.R.C.S., S.E.P. ***Data curation***: C.R.C.S., S.E.P. ***Funding acquisition***: C.R.C.S., S.E.P. ***Investigation***: C.R.C.S. ***Methodology***: C.R.C.S., A.G., R.J.B., S.E.P. ***Project administration***: C.R.C.S., S.E.P. ***Software***: C.R.C.S., A.G. ***Supervision***: S.E.P. ***Visualization***: C.R.C.S., A.G. ***Writing—original draft***: C.R.C.S. ***Writing—review & editing***: C.R.C.S., A.G., R.J.B., S.E.P.

## Data, code, and materials availability

All data, code, and materials needed to evaluate the conclusions in the paper will be provided upon publication alongside the final article. Inquiries regarding data availability should be directed to C.R.C.S. and S.E.P.

## References

1. C. D. N. Barel, Towards a constructional morphology of cichlid fishes (Teleostei, Perciformes). Neth. J. Zool. 33, 357–424 (1983).

2. M. W. Westneat, Transmission of force and velocity in the feeding mechanisms of labrid fishes (Teleostei, Perciformes). Zoomorphology 114, 103–118 (1994).

3. M. W. Westneat, Feeding, function, and phylogeny: analysis of historical biomechanics in labrid fishes using comparative methods. Syst. Biol. 44, 361–383 (1995).

4. M. W. Westneat, A biomechanical model for analysis of muscle force, power output and lower jaw motion in fishes. J. Theor. Biol. 223, 269–281 (2003).

5. M. W. Westneat, Evolution of levers and linkages in the feeding mechanisms of fishes. Integr. Comp. Biol. 44, 378–389 (2004).

6. M. W. Westneat, in Fish Physiology, R. E. Shadwick, G. V. Lauder, Eds. (Academic Press, 2005), vol. 23, chap. 2, pp. 29–75.

7. R. S. Mehta, P. C. Wainwright, Biting releases constraints on moray eel feeding kinematics. J. Exp. Biol. 210, 495–504 (2007).

8. E. Heiss, P. Aerts, S. Van Wassenbergh, Aquatic–terrestrial transitions of feeding systems in vertebrates: a mechanical perspective. J. Exp. Biol. 221, jeb154427 (2018).

9. P. Lemell, N. Natchev, C. J. Beisser, E. Heiss, in Feeding in Vertebrates: Evolution, Morphology, Behavior, Biomechanics, V. Bels, I. Q. Whishaw, Eds. (Springer, 2019), chap. 16, pp. 611–642.

10. D. Schwarz, N. Konow, Y. T. Roba, E. Heiss, A salamander that chews using complex, three-dimensional mandible movements. J. Exp. Biol. 223, jeb220749 (2020).

11. S. M. Gartner et al., Suction feeding of West African lungfish (*Protopterus annectens*): an XROMM analysis of jaw mechanics, cranial kinesis, and hyoid mobility. Biol. Open 11, bio059447 (2022).

12. D. Schwarz, E. Heiss, T. W. Pierson, N. Konow, R. R. Schoch, Using salamanders as model taxa to understand vertebrate feeding constraints during the late Devonian water-to-land transition. Philos. Trans. R. Soc. Lond., Ser. B: Biol. Sci. 378, 20220541 (2023).

13. P. E. Ahlberg, J. A. Clack, Lower jaws, lower tetrapods–a review based on the Devonian genus *Acanthostega*. Trans. R. Soc. Edinb. Earth Sci. 89, 11–46 (1998).

14. P. S. L. Anderson, Biomechanics, functional patterns, and disparity in Late Devonian arthrodires. Paleobiology 35, 321–342 (2009).

15. B. Igielman et al., The lower jaw of Devonian ray-finned fishes (Actinopterygii): anatomy, relationships, and functional morphology. Anat. Rec. 309, 550–602 (2026).

16. L. B. Porro, E. J. Rayfield, J. A. Clack, Descriptive anatomy and three-dimensional reconstruction of the skull of the early tetrapod *Acanthostega gunnari* Jarvik, 1952. PLoS ONE 10, e0118882 (2015).

17. J. B. Lemberg, N. H. Shubin, M. W. Westneat, Feeding kinematics and morphology of the alligator gar (*Atractosteus spatula*, Lacépède, 1803). J. Morphol. 280, 1548–1570 (2019).

18. G. V. Lauder, in Functional Vertebrate Morphology, M. Hildebrand, D. M. Bramble, K. F. Liem, D. B. Wake, Eds. (Harvard University Press, Cambridge and London, 1985), chap. 12, pp. 210–229.

19. G. V. Lauder, H. B. Shaffer, Functional morphology of the feeding mechanism in aquatic ambystomatid salamanders. J. Morphol. 185, 297–326 (1985).

20. W. E. Bemis, G. V. Lauder, Morphology and function of the feeding apparatus of the lungfish, *Lepidosiren paradoxa* (Dipnoi). J. Morphol. 187, 81–108 (1986).

21. G. V. Lauder, in Functional Morphology in Vertebrate Paleontology, J. J. Thomason, Ed. (Cambridge University Press, Cambridge, 1995).

22. A. P. Summers, K. F. Darouian, A. M. Richmond, E. L. Brainerd, Kinematics of aquatic and terrestrial prey capture in *Terrapene carolina*, with implications for the evolution of feeding in cryptodire turtles. J. Exp. Zool. 281, 280–287 (1998).

23. M. E. Alfaro, J. Janovetz, M. W. Westneat, Motor control across trophic strategies: muscle activity of biting and suction feeding fishes. Am. Zool. 41, 1266–1279 (2001).

24. S. M. Deban, S. B. Marks, Metamorphosis and evolution of feeding behaviour in salamanders of the family Plethodontidae. Zool. J. Linn. Soc. 134, 375–400 (2002).

25. A. L. Camp, E. L. Brainerd, Role of axial muscles in powering mouth expansion during suction feeding in largemouth bass (*Micropterus salmoides*). J. Exp. Biol. 217, 1333–1345 (2014).

26. C. M. Stinson, S. M. Deban, Functional trade-offs in the aquatic feeding performance of salamanders. Zoology 125, 69–78 (2017).

27. K. R. Whitlow, C. F. Ross, N. J. Gidmark, J. D. Laurence-Chasen, M. W. Westneat, Suction feeding biomechanics of *Polypterus bichir*: investigating linkage mechanisms and the contributions of cranial kinesis to oral cavity volume change. J. Exp. Biol. 225, jeb243283 (2022).

28. G. Jorgewich-Cohen et al., Morphological diversity of turtle hyoid apparatus is linked to feeding behavior. Integrative Organismal Biology 6, obae014 (2024).

29. T. L. Stubbs, S. E. Pierce, E. J. Rayfield, P. S. L. Anderson, Morphological and biomechanical disparity of crocodile-line archosaurs following the end-Triassic extinction. Proc. R. Soc. Lond., Ser. B: Biol. Sci. 280, 20131940 (2013).

30. D. R. Bellwood, Origins and escalation of herbivory in fishes: a functional perspective. Paleobiology 29, 71–83 (2003).

31. P. S. L. Anderson, M. Friedman, M. D. Brazeau, E. J. Rayfield, Initial radiation of jaws demonstrated stability despite faunal and environmental change. Nature 476, 206–209 (2011).

32. P. S. L. Anderson, M. Friedman, M. Ruta, Late to the table: diversification of tetrapod mandibular biomechanics lagged behind the evolution of terrestriality. Integr. Comp. Biol. 53, 197–208 (2013).

33. J. M. Neenan, M. Ruta, J. A. Clack, E. J. Rayfield, Feeding biomechanics in Acanthostega and across the fish–tetrapod transition. Proc. R. Soc. Lond., Ser. B: Biol. Sci. 281, 20132689 (2014).

34. J. R. G. Rawson et al., Widespread convergence towards functional optimization in the lower jaws of crocodile-line archosaurs. *Proceedings of the Royal Society of London*, Series B 291, 20240720 (2024).

35. J. J. Hill, M. N. Puttick, T. L. Stubbs, E. J. Rayfield, P. C. J. Donoghue, Evolution of jaw disparity in fishes. Palaeontology 61, 847–854 (2018).

36. W. J. Deakin et al., Increasing morphological disparity and decreasing optimality for jaw speed and strength during the radiation of jawed vertebrates. Sci. Adv. 8, eabl3644 (2022).

37. E. M. Troyer et al., Macroevolutionary role reversals in the earliest radiation of bony fishes. Curr. Biol. 35, 4631–4641 (2025).

38. J. Bland et al., Comparison of diverse mandibular mechanics during biting in Devonian lungfishes. iScience 28, 112970 (2025).

39. A. Knapp et al., Adaptive radiation of pelagiarian fishes at the K/Pg boundary led to rapid diversification of mandible morphology. Evol. Lett. 10, 1–11 (2026).

40. P. C. Wainwright, B. A. Richard, Predicting patterns of prey use from morphology of fishes. Environ. Biol. Fishes 44, 97–113 (1995).

41. C. D. Hulsey, M. C. Mims, N. F. Parnell, J. T. Streelman, Comparative rates of lower jaw diversification in cichlid adaptive radiations. J. Evol. Biol. 23, 1456–1467 (2010).

42. S. A. Price, R. Holzman, T. J. Near, P. C. Wainwright, Coral reefs promote the evolution of morphological diversity and ecological novelty in labrid fishes. Ecol. Lett. 14, 462–469 (2011).

43. R. Holzman et al., Biomechanical trade-offs bias rates of evolution in the feeding apparatus of fishes. *Proceedings of the Royal Society of London*, Series B 279, 1287–1292 (2012).

44. H. O. Berks, P. S. M. Carmona, P. C. J. Donoghue, E. J. Rayfield, The evolution of herbivory, not terrestrialization, drove morphological change in the mandibles of Palaeozoic tetrapods. Evol. J. Linn. Soc. 4, kzaf004 (2025).

45. J. Ponstein, M. J. MacDougall, J. Schaeffer, C. F. Kammerer, J. Fröbisch, Mandibular form and function is more disparate in amniotes than in non-amniote tetrapods from the late Palaeozoic. PeerJ 13, e20243 (2025).

46. M. L. Zelditch, J. Ye, J. S. Mitchell, D. L. Swiderski, Rare ecomorphological convergence on a complex adaptive landscape: body size and diet mediate evolution of jaw shape in squirrels (Sciuridae). Evolution 71, 633–649 (2017).

47. D. M. Grossnickle, Feeding ecology has a stronger evolutionary influence on functional morphology than on body mass in mammals. Evolution 74, 610–628 (2020).

48. N. M. Morales-García, P. G. Gill, C. M. Janis, E. J. Rayfield, Jaw shape and mechanical advantage are indicative of diet in Mesozoic mammals. *Commun*. Biol. 4, 242 (2021).

49. D. L. Swiderski, M. L. Zelditch, Complex adaptive landscape for a “simple” structure: the role of trade-offs in the evolutionary dynamics of mandibular shape in ground squirrels. Evolution 76, 946–965 (2022).

50. Z. J. Tseng et al., A switch in jaw form–function coupling during the evolution of mammals. Philos. Trans. R. Soc. Lond., Ser. B: Biol. Sci. 378, 20220091 (2023).

51. B. W. Griffin et al., Evolution of chondrichthyan jaw morphology, from ecological generalists to specialists. Paleobiology 52, 131–144 (2026).

52. M. A. MacIver, L. Schmitz, U. Mugan, T. D. Murphey, C. D. Mobley, Massive increase in visual range preceded the origin of terrestrial vertebrates. Proc. Natl. Acad. Sci. 114, E2375–E2384 (2017).

53. J. A. Clack, Devonian climate change, breathing, and the origin of the tetrapod stem group. Integr. Comp. Biol. 47, 510–523 (2007).

54. J. B. Lemberg, E. B. Daeschler, N. H. Shubin, The feeding system of Tiktaalik roseae: an intermediate between suction feeding and biting. Proc. Natl. Acad. Sci. 118, e2016421118 (2021).

55. E. L. Brainerd, Gradual evolution of costal aspiration breathing in tetrapods. Anat. Rec. 309, 1631–1643 (2026).

56. J. A. Clack, Patterns and processes in the early evolution of the tetrapod ear. J. Neurobiol. 53, 251–264 (2002).

57. B. V. Dickson, J. A. Clack, T. R. Smithson, S. E. Pierce, Functional adaptive landscapes predict terrestrial capacity at the origin of limbs. Nature 589, 242–245 (2021).

58. S. E. Pierce, J. R. Hutchinson, J. A. Clack, Historical perspectives on the evolution of tetrapodomorph movement. Integr. Comp. Biol. 53, 209–223 (2013).

59. M. J. Markey, C. R. Marshall, Terrestrial-style feeding in a very early aquatic tetrapod is supported by evidence from experimental analysis of suture morphology. Proc. Natl. Acad. Sci. 104, 7134–7138 (2007).

60. L. B. Porro, E. J. Rayfield, J. A. Clack, Computed tomography, anatomical description and three-dimensional reconstruction of the lower jaw of *Eusthenopteron foordi* Whiteaves, 1881 from the Upper Devonian of Canada. Palaeontology 58, 1031–1047 (2015).

61. J. R. G. Rawson, L. B. Porro, E. Martin-Silverstone, E. J. Rayfield, Osteology and digital reconstruction of the skull of the early tetrapod Whatcheeria deltae. J. Vert. Paleontol. 41, e1927749 (2021).

62. J. R. G. Rawson, B. Esteve-Altava, L. B. Porro, H. Dutel, E. J. Rayfield, Early tetrapod cranial evolution is characterized by increased complexity, constraint, and an offset from fin-limb evolution. Sci. Adv. 8, eadc8875 (2022).

63. L. B. Porro, E. Martin-Silverstone, E. J. Rayfield, Descriptive anatomy and three-dimensional reconstruction of the skull of the tetrapod *Eoherpeton watsoni* Panchen, 1975 from the Carboniferous of Scotland. Earth Environ. Sci. Trans. Royal Soc. Edinburgh 115, 164–184 (2024).

64. L. B. Porro, E. J. Rayfield, J. A. Clack, Computed tomography and three-dimensional reconstruction of the skull of the stem tetrapod *Crassigyrinus scoticus* Watson, 1929. J. Vert. Paleontol. 42, e2183134 (2022).

65. M. E. Alfaro, D. I. Bolnick, P. C. Wainwright, Evolutionary dynamics of complex biomechanical systems: an example using the four-bar mechanism. Evolution 58, 495–503 (2004).

66. P. C. Wainwright, M. E. Alfaro, D. I. Bolnick, C. D. Hulsey, Many-to-one mapping of form to function: a general principle in organismal design? Integr. Comp. Biol. 45, 256–262 (2005).

67. G. Sansalone, S. Wroe, G. Coates, M. R. G. Attard, C. Fruciano, Unexpectedly uneven distribution of functional trade-offs explains cranial morphological diversity in carnivores. Nat. Commun. 15, 3275 (2024).

68. M. E. Alfaro, D. I. Bolnick, P. C. Wainwright, Evolutionary consequences of many-to-one mapping of jaw morphology to mechanics in labrid fishes. The American Naturalist 165, E140–E154 (2005).

69. J. P. Friel, P. C. Wainwright, Evolution of motor patterns in tetraodontiform fishes: does muscle duplication lead to functional diversification? Brain Behav. Evol. 52, 159–170 (1998).

70. E. G. Axlid, S. Shirazi, T. E. Higham, Feeding in flow: environmental flow conditions alter prey capture kinematics in fish. Ecol. Evol. Physiol. 0, 1–33 (2026).

71. E. D. Burress, C. M. Martinez, P. C. Wainwright, Decoupled jaws promote trophic diversity in cichlid fishes. Evolution 74, 950–961 (2020).

72. A. Velasco-Hogan, W. Huang, C. Serrano, D. Kisailus, M. A. Meyers, Tooth structure, mechanical properties, and diet specialization of Piranha and Pacu (Serrasalmidae): a comparative study. Acta Biomater. 134, 531–545 (2021).

73. C. D. Marshall et al., The ontogenetic scaling of bite force and head size in loggerhead sea turtles (*Caretta caretta*): implications for durophagy in neritic, benthic habitats. J. Exp. Biol. 215, 4166–4174 (2012).

74. R. Holzman, D. C. Collar, R. S. Mehta, P. C. Wainwright, Functional complexity can mitigate performance trade-offs. The American Naturalist 177, E69–E83 (2011).

75. J. M. Gardiner, P. J. Motta, Largemouth bass (*Micropterus salmoides*) switch feeding modalities in response to sensory deprivation. Zoology 115, 78–83 (2012).

76. C. D. Wilga, P. J. Motta, C. P. Sanford, Evolution and ecology of feeding in elasmobranchs. Integr. Comp. Biol. 47, 55–69 (2007).

77. R. Holmes, The skull and axial skeleton of the Lower Permian anthracosauroid amphibian *Archeria crassidisca* Cope. Palaeontographica 207, 161–206 (1989).

78. T. Skorczewski, A. Cheer, P. C. Wainwright, The benefits of planar circular mouths on suction feeding performance. J. R. Soc. Interface 9, 1767–1773 (2012).

79. J. M. Huie, A. P. Summers, M. A. Kolmann, Body shape separates guilds of rheophilic herbivores (Myleinae: Serrasalmidae) better than feeding morphology. Proc. Acad. Nat. Sci. Phila. 166, 1–15 (2020).

80. L. Leuzinger, L. Cavin, A. López-Arbarello, J.-P. Billon-Bruyat, Peculiar tooth renewal in a Jurassic ray-finned fish (Lepisosteiformes, †Scheenstia sp.). Palaeontology 63, 117–129 (2020).

81. K. E. Cohen, O. Lucanus, A. P. Summers, M. A. Kolmann, Lip service: histological phenotypes correlate with diet and feeding ecology in herbivorous pacus. Anat. Rec. 306, 326–342 (2023).

82. D. C. Collar, P. C. Wainwright, M. E. Alfaro, L. J. Revell, R. S. Mehta, Biting disrupts integration to spur skull evolution in eels. Nat. Commun. 5, 5505 (2014).

83. C. Meloro, D. Tamagnini, Macroevolutionary ecomorphology of the Carnivora skull: adaptations and constraints in the extant species. Zool. J. Linn. Soc. 196, 1054–1068 (2022).

84. D. H. Nemeth, Modulation of attack behavior and its effect on feeding performance in a trophic generalist fish, *Hexagrammos decagrammus*. J. Exp. Biol. 200, 2155–2164 (1997).

85. A. Huby et al., Functional diversity in biters: the evolutionary morphology of the oral jaw system in pacus, piranhas and relatives (Teleostei: Serrasalmidae). Biol. J. Linn. Soc. 127, 722–741 (2019).

86. C. E. Oufiero, R. A. Holzman, F. A. Young, P. C. Wainwright, New insights from serranid fishes on the role of trade-offs in suction-feeding diversification. J. Exp. Biol. 215, 3845–3855 (2012).

87. M. J. McHenry, There is no trade-off between speed and force in a dynamic lever system. Biol. Lett. 7, 384–386 (2011).

88. C. N. Jacobs, R. Holzman, Conserved spatio-temporal patterns of suction-feeding flows across aquatic vertebrates: a comparative flow visualization study. J. Exp. Biol. 221, jeb174912 (2018).

89. K. F. Liem, in Trophic Interactions Within Aquatic Ecosystems, D. G. Meyers, J. R. Strickler, Eds. (Routledge, 1984), pp. 269–305.

90. G. G. Sass, P. J. Motta, The effects of satiation on strike mode and prey capture kinematics in the largemouth bass, *Micropterus salmoides*. Environ. Biol. Fishes 65, 441–454 (2002).

91. S. J. Longo, M. D. McGee, C. E. Oufiero, T. B. Waltzek, P. C. Wainwright, Body ram, not suction, is the primary axis of suction-feeding diversity in spiny-rayed fishes. J. Exp. Biol. 219, 119–128 (2016).

92. M. W. Caldwell, The Origin of Snakes: Morphology and the Fossil Record. (Taylor & Francis, Boca Raton, 2019).

93. J. Davenport, M. Spikes, S. M. Thornton, B. O. Kelly, Crab-eating in the diamondback terrapin *Malaclemys terrapin*: dealing with dangerous prey. J. Mar. Biol. Assoc. U.K. 72, 835–848 (1992).

94. V. L. Bels, J. Davenport, S. Renous, Food ingestion in the estuarine turtle *Malaclemys terrapin*: comparison with the marine leatherback turtle *Dermochelys coriacea*. J. Mar. Biol. Assoc. U.K. 78, 953–972 (1998).

95. N. Natchev, E. Heiss, P. Lemell, D. Stratev, J. Weisgram, Analysis of prey capture and food transport kinematics in two Asian box turtles, *Cuora amboinensis* and *Cuora flavomarginata* (Chelonia, Geoemydidae), with emphasis on terrestrial feeding patterns. Zoology 112, 113–127 (2009).

96. M. E. H. Jones et al., The head and neck anatomy of sea turtles (Cryptodira: Chelonioidea) and skull shape in Testudines. PLoS ONE 7, e47852 (2012).

97. A. Guzman, Texas A&M University, (2008).

98. N. Natchev, N. Tzankov, I. Werneburg, E. Heiss, Feeding behaviour in a ‘basal’ tortoise provides insights on the transitional feeding mode at the dawn of modern land turtle evolution. PeerJ 3, e1172 (2015).

99. L. W. Witmer, in Functional Morphology in Vertebrate Paleontology, J. Thomason, Ed. (Cambridge University Press, 1995), pp. 19–33.

100. S. Van Wassenbergh, A. Herrel, D. Adriaens, P. Aerts, No trade-off between biting and suction feeding performance in clariid catfishes. J. Exp. Biol. 210, 27–36 (2007).

101. O. Shoval et al., Evolutionary trade-offs, Pareto optimality, and the geometry of phenotype space. Science 336, 1157–1160 (2012).

102. G. V. Lauder, H. B. Shaffer, Functional design of the feeding mechanism in lower vertebrates: unidirectional and bidirectional flow systems in the tiger salamander. Zool. J. Linn. Soc. 88, 277–290 (1986).

103. G. V. Lauder, H. B. Shaffer, Ontogeny of functional design in tiger salamanders (*Ambystoma tigrinum*): are motor patterns conserved during major morphological transformations? J. Morphol. 197, 249–268 (1988).

104. S. M. Reilly, G. V. Lauder, Ontogeny of aquatic feeding performance in the eastern newt, *Notophthalmus viridescens* (Salamandridae). Copeia 1988, 87–91 (1988).

105. W. E. Bemis, Morphology and growth of lepidosirenid lungfish tooth plates (Pisces: Dipnoi). J. Morphol. 179, 73–93 (1984).

106. W. E. Bemis, Feeding systems of living Dipnoi: anatomy and function. J. Morphol. 190, 249–275 (1986).

107. D. Schwarz, M. T. Fedler, P. Lukas, A. Kupfer, Form and function of the feeding apparatus of sirenid salamanders (Caudata: Sirenidae): three-dimensional chewing and herbivory? Zool. Anz. 295, 99–116 (2021).

108. B. T. Miller, J. H. Larsen, Jr, Feeding performance in aquatic postmetamorphic newts (Urodela: Salamandridae): are bidirectional flow systems necessarily inefficient? Canadian Journal of Zoology 67, 2414–2421 (1989).

109. J. T. Streelman, M. Alfaro, M. W. Westneat, D. R. Bellwood, S. A. Karl, Evolutionary history of the parrotfishes: biogeography, ecomorphology, and comparative diversity. Evolution 56, 961–971 (2002).

110. R. S. Mehta, P. C. Wainwright, Raptorial jaws in the throat help moray eels swallow large prey. Nature 449, 79–82 (2007).

111. V. L. Bels, J. Davenport, V. Delheusy, Kinematic analysis of the feeding behavior in the box turtle *Terrapene carolina* (L.), (Reptilia: Emydidae). J. Exp. Zool. 277, 198–212 (1997).

112. E. H. Beaumont, T. R. Smithson, The cranial morphology and relationships of the aberrant Carboniferous amphibian *Spathicephalus mirus* Watson. Zool. J. Linn. Soc. 122, 187–209 (1998).

113. J. A. Clack et al., *Acherontiscus caledoniae*: the earliest heterodont and durophagous tetrapod. R. Soc. Open Sci. 6, 182087 (2019).

114. T. R. Simões, S. E. Pierce, Sustained high rates of morphological evolution during the rise of tetrapods. *Nat*. Ecol. Evol. 5, 1403–1414 (2021).

115. S. Kumar et al., TimeTree 5: an expanded resource for species divergence times. Mol. Biol. Evol. 39, msac174 (2022).

116. R Core Team. (R Foundation for Statistical Computing, Vienna, Austria, 2023).

117. RStudio Team. (RStudio, PBC, Boston, MA, 2023).

118. D. W. Bapst, paleotree: an R package for paleontological and phylogenetic analyses of evolution. Methods Ecol. Evol. 3, 803–807 (2012).

119. J. Alroy, C. Marshall, A. Miller. (2004).

120. Materialise. (Belgium, 2021–2024).

121. Epic Games Slovakia. (Slovakia, 2025).

122. P. Cignoni et al., paper presented at the Sixth Eurographics Italian Chapter Conference, 2008.

123. P. Cignoni et al. (2022).

124. Agisoft. (Agisoft LLC, 2018).

125. CloudCompare. (2024).

126. Materialise. (Belgium, 2021–2025).

127. V. Bonhomme, S. Picq, C. Gaucherel, J. Claude, Momocs: outline analysis using R. Journal of Statistical Software 56, 1–24 (2014).

128. F. J. Rohlf, The tps series of software. *Hystrix Ital*. J. Mammal. 26, 9–12 (2015).

129. F. J. Rohlf. (Department of Ecology and Evolution, State University of New York at Stony Brook, 2021).

130. F. J. Rohlf. (Department of Ecology and Evolution, State University of New York at Stony Brook, 2023).

131. Robert McNeel & Associates. (2024).

132. Robert McNeel & Associates. (2024).

133. H. Wickham. (Springer-Verlag, New York, 2016).

134. I. Patil, Visualizations with statistical details: the ‘ggstatsplot’ approach. Journal of Open Source Software 6, 3167 (2021).

135. C. A. Martin. (2017).

136. J. Oksanen et al. (2024).

137. P. Martinez Arbizu. (2017).

138. C. R. John, D. Watson, M. R. Barnes, C. Pitzalis, M. J. Lewis, Spectrum: fast density-aware spectral clustering for single and multi-omic data. Bioinformatics 36, 1159–1166 (2020).

139. C. R. John, D. Watson. (2020).

140. A. Gillet, K. E. Jones, S. E. Pierce, Repatterning of mammalian backbone regionalization in cetaceans. Nat. Commun. 15, 7587 (2024).

141. C. Weihs, U. Ligges, K. Luebke, N. Raabe, in Data Analysis and Decision Support, D. Baier, R. Decker, L. Schmidt-Thieme, Eds. (Springer-Verlag, Berlin, 2005), pp. 335–343.

142. M. Friendly, HE plots for multivariate linear models. J. Comput. Graph. Stat. 16, 421–444 (2007).

143. M. Friendly, J. Fox. (2024).

144. J. C. Pinheiro, D. M. Bates, Mixed-Effects Models in S and S-PLUS. (Springer, New York, 2000).

145. J. C. Pinheiro, D. M. Bates, R Core Team. (2023).

146. E. Paradis, J. Claude, K. Strimmer, APE: Analyses of Phylogenetics and Evolution in R language. Bioinformatics 20, 289–290 (2004).

147. E. Paradis, K. Schliep, ape 5.0: an environment for modern phylogenetic and evolutionary analyses in R. Bioinformatics 35, 526–528 (2019).

