## Supplementary Material for "Playing by their own rules: clade-specific trends in mandibular form and function disrupt current paradigms of early tetrapod evolution"

### Supplementary Materials

#### Supplementary Tables

The following supplementary tables accompany this manuscript, and will be provided upon publication (see Data Availability statement for details).

Table S1. **Taxon sample and data**.

Table S2. **Coordinates from principal components analysis of mandibular functional metrics**.

Table S3. **Allometric scaling of principal components scores *versus* mandible size**.

Table S4. **Coordinates, predicted feeding modes, and posterior probabilities from linear discriminant analysis of mandibular functional metrics, under a 3-mode feeding classification scheme**.

Table S5. **Coordinates, predicted feeding modes, and posterior probabilities from linear discriminant analysis of mandibular functional metrics, under a 5-mode feeding classification scheme**.

Table S6. **Coordinates and cluster assignments from principal components analysis and spectral clustering analysis, respectively, of mandible silhouettes**.

#### Supplementary Data

The following supplementary data files accompany this manuscript, and will be provided upon publication (see Data Availability statement for details).

Data S1. **Phylogeny**.

Data S2. **Source phylogenies and code**.

Data S3. **Mandible silhouettes and landmarks**.

Data S4. **Rhino/Grasshopper macro**.

Data S5. **R scripts**.

#### Supplementary Figures

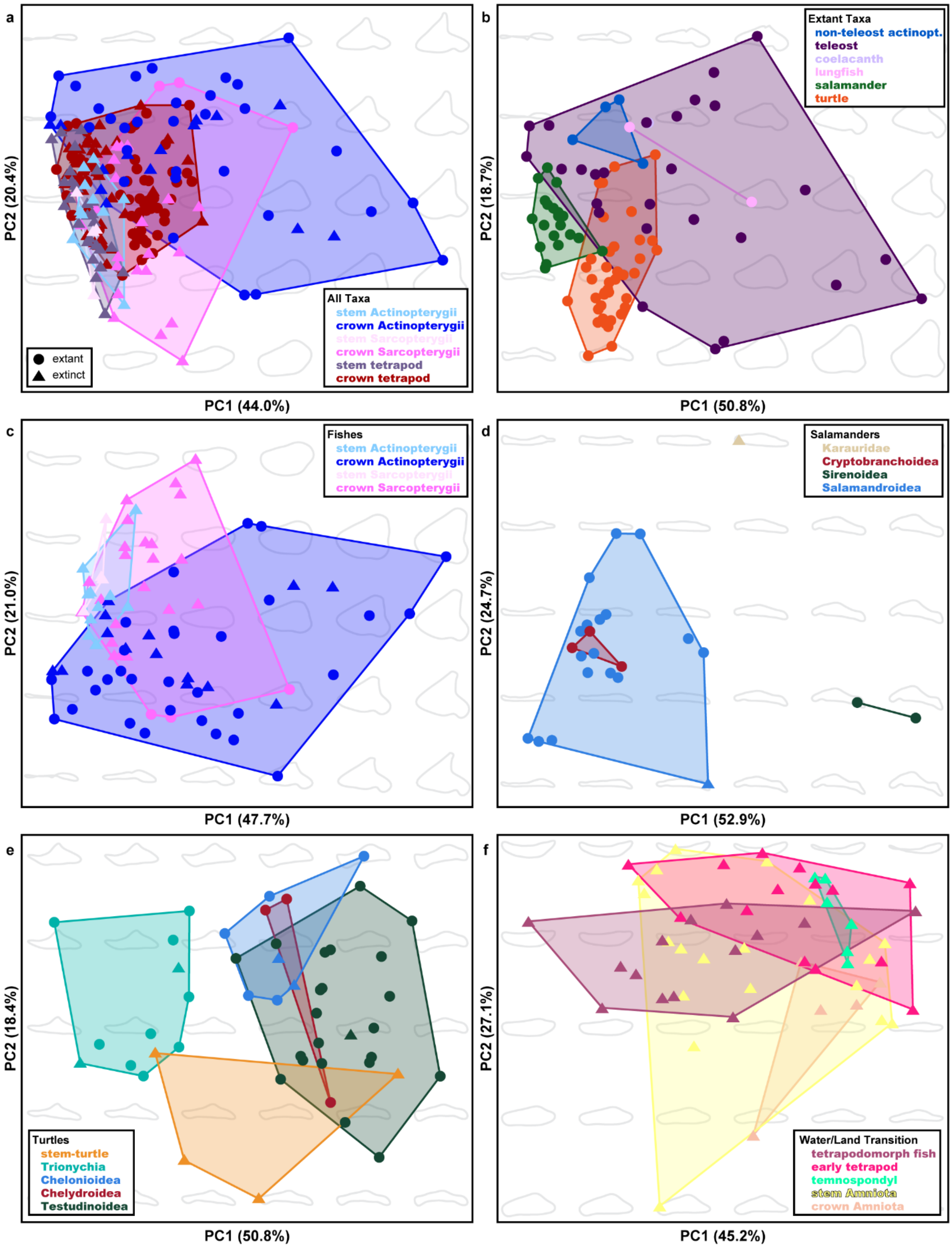
a
PC2 (20.4%)
PC1 (44.0%)
extant
extinct
All Taxa
stem Actinopterygii
crown Actinopterygii
stem Sarcopterygii
crown Sarcopterygii
stem tetrapod
crown tetrapod
b
PC2 (18.7%)
PC1 (50.8%)
Extant Taxa
non-teleost actinopt.
teleost
coelacanth
lungfish
salamander
turtle
c
PC2 (21.0%)
PC1 (47.7%)
Fishes
stem Actinopterygii
crown Actinopterygii
stem Sarcopterygii
crown Sarcopterygii
d
PC2 (24.7%)
PC1 (52.9%)
Salamanders
Karauridae
Cryptobranchoidea
Sirenoidea
Salamandroidea
e
PC2 (18.4%)
PC1 (50.8%)
Turtles
stem-turtle
Trionychia
Chelonioidea
Chelydroidea
Testudinoidea
f
PC2 (27.1%)
PC1 (45.2%)
Water/Land Transition
tetrapodomorph fish
early tetrapod
temnospondyl
stem Amniota
crown Amniota

Figure S1. **Morphospace for key taxon partitions, reflecting principal components analysis of mandible silhouettes, colour-coded by higher taxon**. (**a**) All taxa (n = 214). (**b**) Extant taxa (n = 95). (**c**) Fishes (n = 84). (**d**) Salamanders (n = 24). (**e**) Turtles (n = 48). (**f**) Taxa spanning the tetrapod water-to-land transition (n = 58). Colours indicate taxonomic grouping; see respective legend for each panel. Shape denotes status as extant or extinct; see legend in (a) for shapes used throughout all panels.

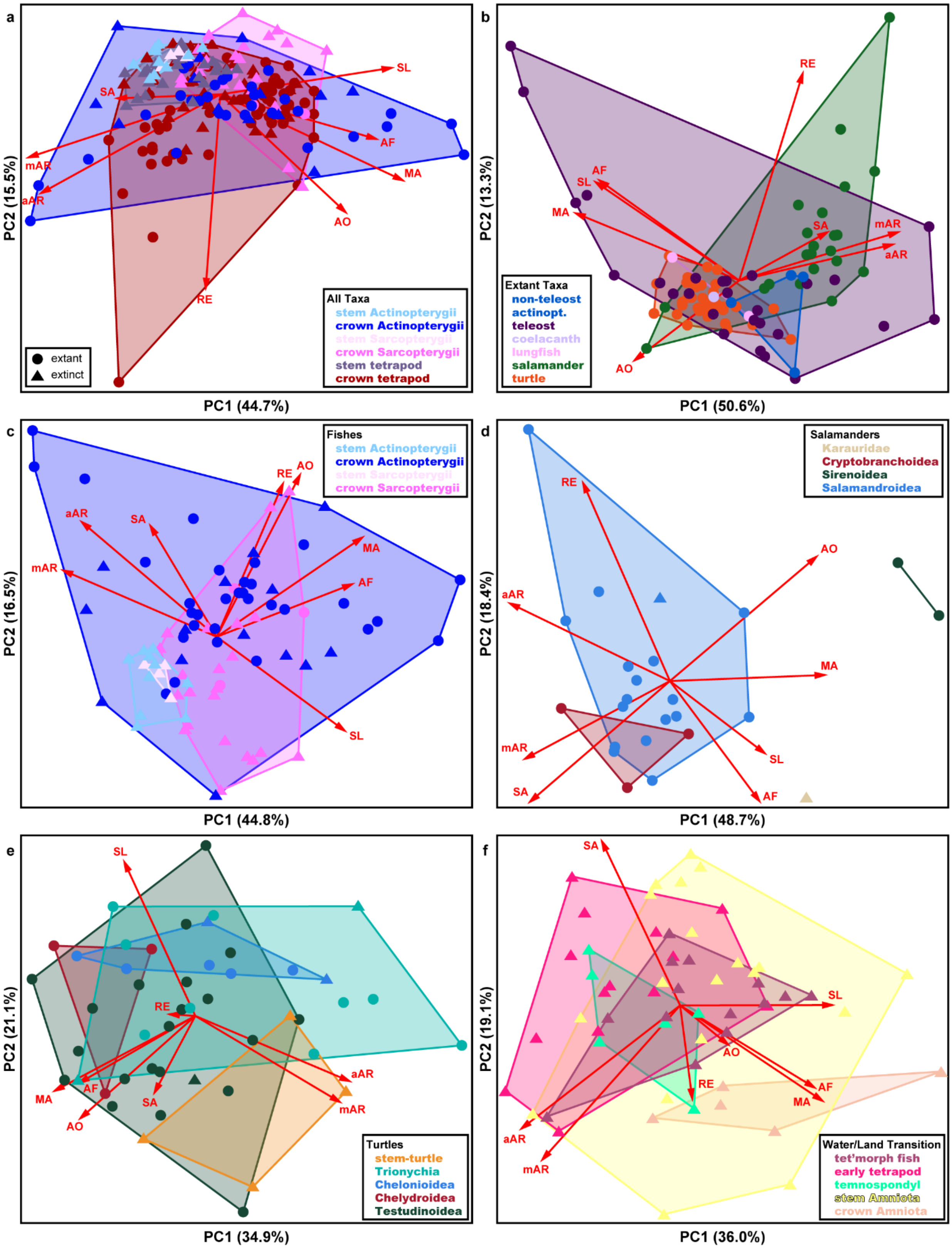
a
PC1 (44.7%)
PC2 (15.5%)
SL
SA
mAR
aAR
AF
MA
AO
RE
extant
extinct
All Taxa
stem Actinopterygii
crown Actinopterygii
stem Sarcopterygii
crown Sarcopterygii
stem tetrapod
crown tetrapod
b
PC1 (50.6%)
PC2 (13.3%)
RE
AF
SL
MA
SA
mAR
aAR
AO
Extant Taxa
non-teleost actinopt.
teleost
coelacanth
lungfish
salamander
turtle
c
PC1 (44.8%)
PC2 (16.5%)
RE
AO
aAR
SA
MA
mAR
AF
SL
Fishes
stem Actinopterygii
crown Actinopterygii
stem Sarcopterygii
crown Sarcopterygii
d
PC1 (48.7%)
PC2 (18.4%)
RE
AO
aAR
MA
mAR
SL
SA
AF
Salamanders
Karauridae
Cryptobranchoidea
Sirenoidea
Salamandroidea
e
PC1 (34.9%)
PC2 (21.1%)
SL
RE
MA
AF
SA
AO
aAR
mAR
Turtles
stem-turtle
Trionychia
Chelonioidea
Chelydroidea
Testudinoidea
f
PC1 (36.0%)
PC2 (19.1%)
SA
SL
AO
RE
AF
MA
aAR
mAR
Water/Land Transition
tet'morph fish
early tetrapod
temnospondyl
stem Amniota
crown Amniota

Figure S2. **Mechanospace for key taxon partitions, reflecting principal components analysis of functional metrics, colour-coded by higher taxon**. (**a**) All taxa (n = 214). (**b**) Extant taxa (n = 95). (**c**) Fishes (n = 84). (**d**) Salamanders (n = 24). (**e**) Turtles (n = 48). (**f**) Taxa spanning the tetrapod water-to-land transition (n = 58). Colours indicate taxonomic grouping; see respective legend for each panel. Shape denotes status as extant or extinct; see legend in (a) for shapes used throughout all panels. Biplots indicate the respective contribution of each functional metric to the first two principal components. See Methods for abbreviations of functional metrics.

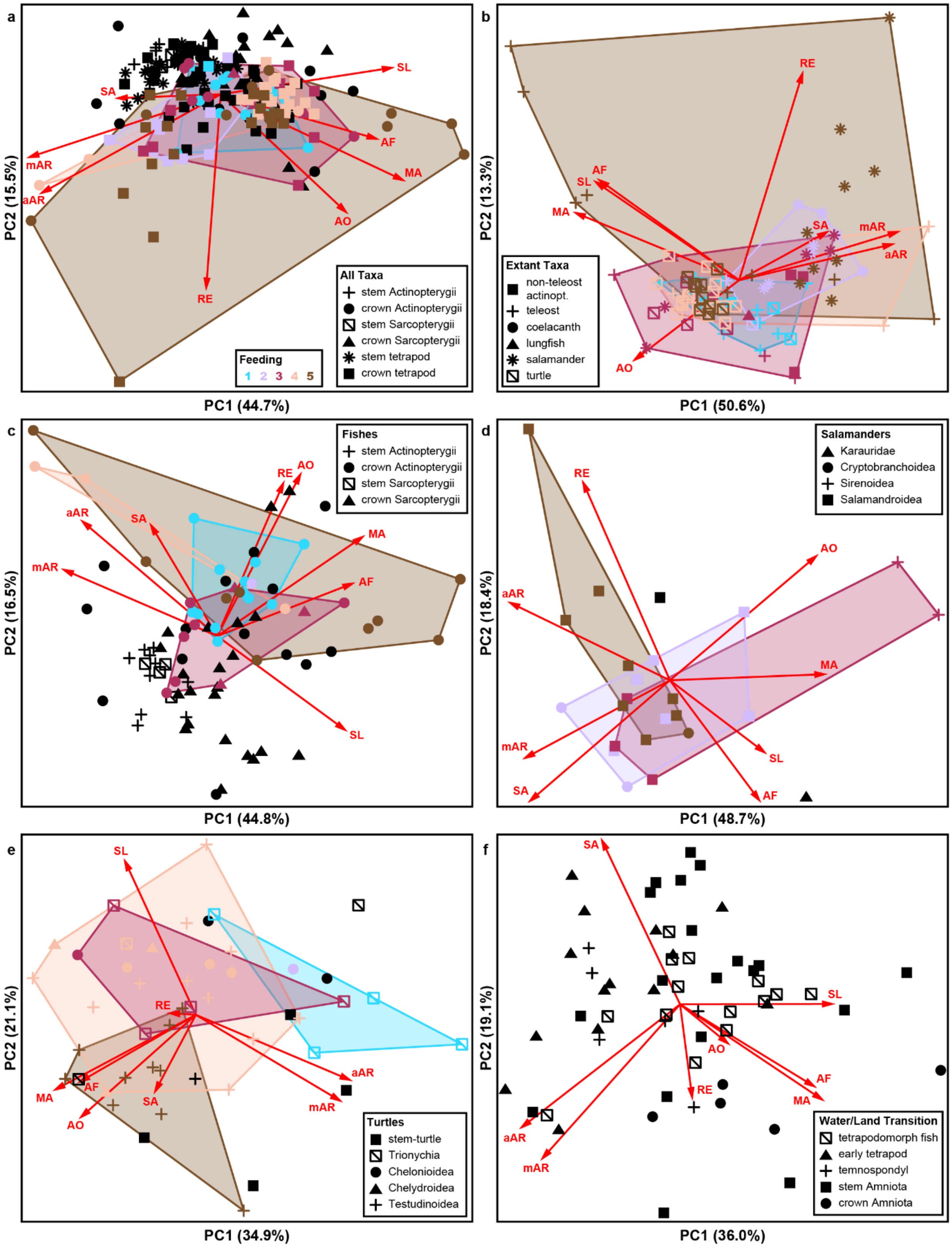
a
SL
SA
mAR
aAR
AF
MA
AO
RE
PC2 (15.5%)
PC1 (44.7%)
Feeding
1 2 3 4 5
All Taxa
stem Actinopterygii
crown Actinopterygii
stem Sarcopterygii
crown Sarcopterygii
stem tetrapod
crown tetrapod
b
RE
AF
SL
MA
SA
mAR
aAR
AO
PC2 (13.3%)
PC1 (50.6%)
Extant Taxa
non-teleost actinopt.
teleost
coelacanth
lungfish
salamander
turtle
c
RE
AO
aAR
SA
MA
mAR
AF
SL
PC2 (16.5%)
PC1 (44.8%)
Fishes
stem Actinopterygii
crown Actinopterygii
stem Sarcopterygii
crown Sarcopterygii
d
RE
AO
aAR
MA
mAR
SA
SL
AF
PC2 (18.4%)
PC1 (48.7%)
Salamanders
Karauridae
Cryptobranchoidea
Sirenoidea
Salamandroidea
e
SL
RE
AF
MA
SA
AO
aAR
mAR
PC2 (21.1%)
PC1 (34.9%)
Turtles
stem-turtle
Trionychia
Chelonioidea
Chelydroidea
Testudinoidea
f
SA
SL
AO
RE
AF
MA
aAR
mAR
PC2 (19.1%)
PC1 (36.0%)
Water/Land Transition
tetrapodomorph fish
early tetrapod
temnospondyl
stem Amniota
crown Amniota

Figure S3. **Mechanospace for key taxon partitions, reflecting principal components analysis of functional metrics, colour-coded using a 5-category feeding mode classification**. (**a**) All taxa (n = 214). (**b**) Extant taxa (n = 95). (**c**) Fishes (n = 84). (**d**) Salamanders (n = 24). (**e**) Turtles (n = 48). (**f**) Taxa spanning the tetrapod water-to-land transition (n = 58). Colours indicate feeding modes for extant taxa; see legend in (a) for colour scheme used throughout all panels. Black datapoints denote extinct taxa. Shape denotes higher taxon; see respective legend for each panel. Biplots indicate the respective contribution of each functional metric to the first two principal components. See Methods for abbreviations of functional metrics.

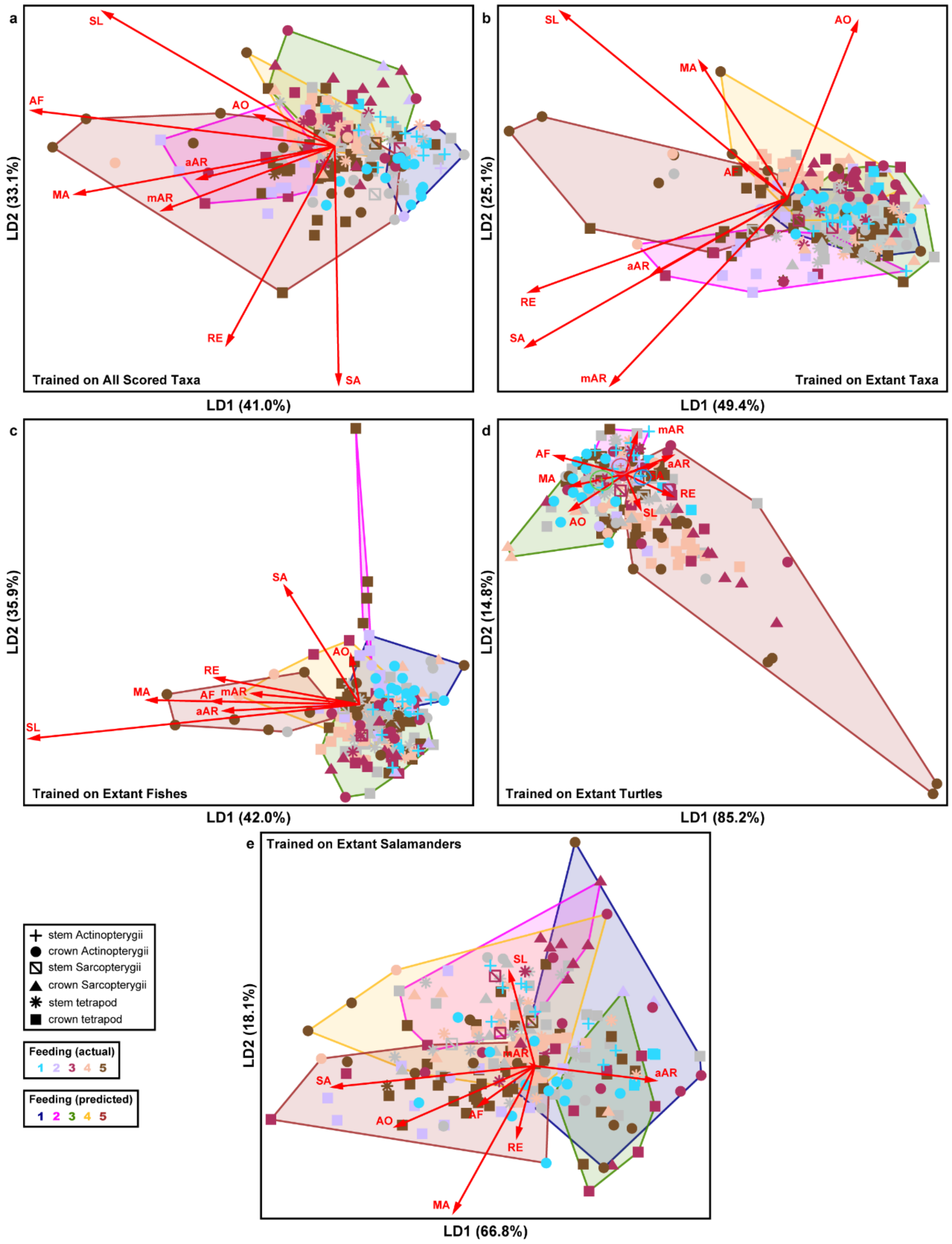
a
b
c
d
e
Trained on All Scored Taxa
Trained on Extant Taxa
Trained on Extant Fishes
Trained on Extant Turtles
Trained on Extant Salamanders
LD1 (41.0%)
LD2 (33.1%)
LD1 (49.4%)
LD2 (25.1%)
LD1 (42.0%)
LD2 (35.9%)
LD1 (85.2%)
LD2 (14.8%)
LD1 (66.8%)
LD2 (18.1%)
SL
AF
AO
aAR
MA
mAR
RE
SA
stem Actinopterygii
crown Actinopterygii
stem Sarcopterygii
crown Sarcopterygii
stem tetrapod
crown tetrapod
Feeding (actual)
1 2 3 4 5
Feeding (predicted)
1 2 3 4 5

Figure S4. **Multivariate space of key taxon partitions, reflecting linear discriminant analysis (LDA) of functional metrics, constructed using a 5-category feeding mode classification**. (**a**) LDA model trained using all taxa for which feeding modes could be ascertained (n = 160, comprising 95 extant and 65 extinct taxa). (**b**) LDA model trained using all extant taxa (n = 95). (**c**) LDA model trained using extant fishes (n = 34). (**d**) LDA model trained using extant salamanders (n = 22). (**e**) LDA model trained using extant turtles (n = 39). Coloured datapoints denote known feeding modes (including among fossils), with grey representing extinct taxa for which feeding mode could not be ascertained from the literature. Coloured hulls reflect feeding modes as predicted by the discriminant model. Shape denotes higher taxon. See legends for colour schemes and shapes used throughout all panels. Biplots indicate the respective loading of each functional metric in the discriminant model. See Methods for abbreviations of functional metrics.

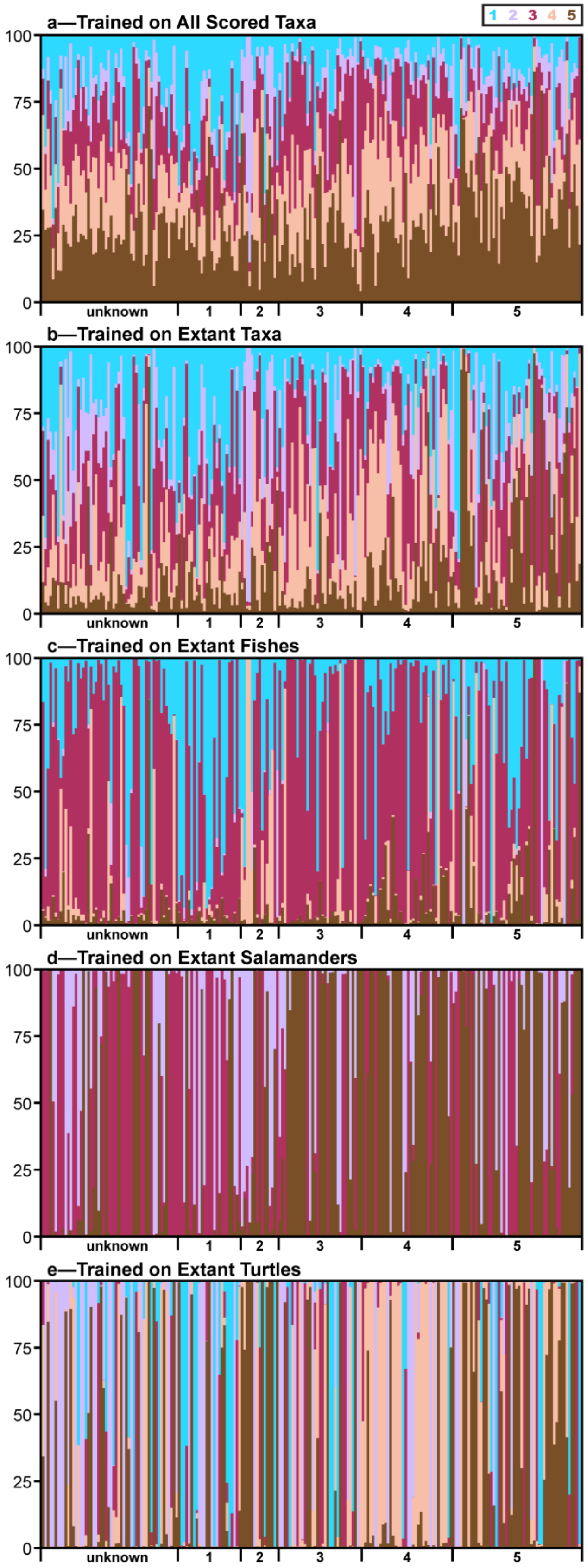
a—Trained on All Scored Taxa
b—Trained on Extant Taxa
c—Trained on Extant Fishes
d—Trained on Extant Salamanders
e—Trained on Extant Turtles
1 2 3 4 5
100
75
50
25
0
unknown
1
2
3
4
5

Figure S5. **Posterior probability of feeding mode classifications, based on linear discriminant analysis (LDA) of key taxon partitions, constructed using a 5-category feeding mode classification**. (**a**) LDA model trained using all taxa for which feeding modes could be ascertained (n = 160, comprising 95 extant and 65 extinct taxa). (**b**) LDA model trained using all extant taxa (n = 95). (**c**) LDA model trained using extant fishes (n = 34). (**d**) LDA model trained using extant salamanders (n = 22). (**e**) LDA model trained using extant turtles (n = 39). Coloured bars represent posterior probability of each taxon belonging to the corresponding feeding mode; see legend at top for colour scheme used throughout all panels.

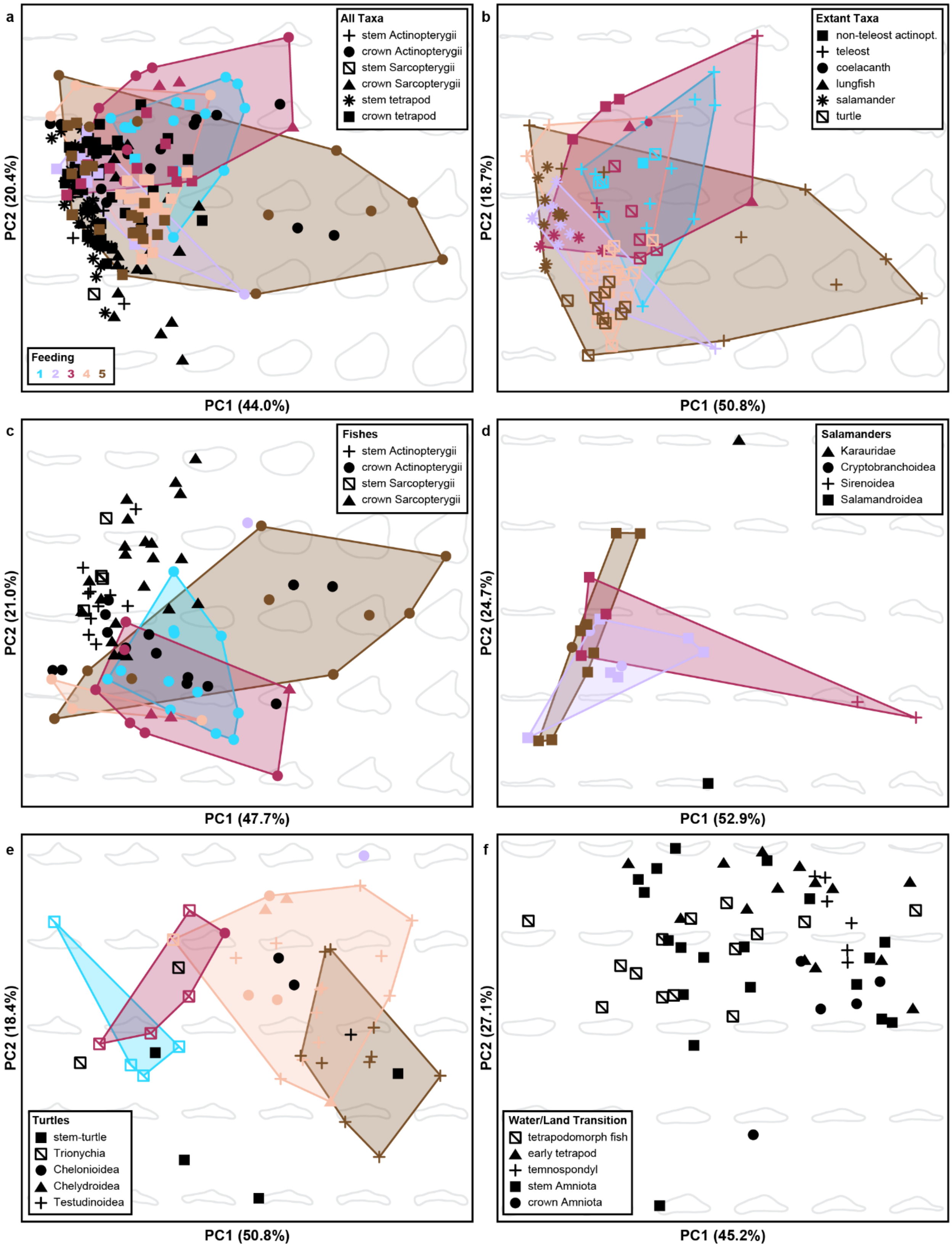
a
All Taxa
stem Actinopterygii
crown Actinopterygii
stem Sarcopterygii
crown Sarcopterygii
stem tetrapod
crown tetrapod
Feeding
1 2 3 4 5
PC2 (20.4%)
PC1 (44.0%)
b
Extant Taxa
non-teleost actinopt.
teleost
coelacanth
lungfish
salamander
turtle
PC2 (18.7%)
PC1 (50.8%)
c
Fishes
stem Actinopterygii
crown Actinopterygii
stem Sarcopterygii
crown Sarcopterygii
PC2 (21.0%)
PC1 (47.7%)
d
Salamanders
Karauridae
Cryptobranchoidea
Sirenoidea
Salamandroidea
PC2 (24.7%)
PC1 (52.9%)
e
Turtles
stem-turtle
Trionychia
Chelonioidea
Chelydroidea
Testudinoidea
PC2 (18.4%)
PC1 (50.8%)
f
Water/Land Transition
tetrapodomorph fish
early tetrapod
temnospondyl
stem Amniota
crown Amniota
PC2 (27.1%)
PC1 (45.2%)

Figure S6. **Morphospace for key taxon partitions, reflecting principal components analysis of mandible silhouettes, colour-coded using a 5-category feeding mode classification**. (**a**) All taxa (n = 214). (**b**) Extant taxa (n = 95). (**c**) Fishes (n = 84). (**d**) Salamanders (n = 24). (**e**) Turtles (n = 48). (**f**) Taxa spanning the tetrapod water-to-land transition (n = 58). Colours indicate feeding modes for extant taxa; see legend in (a) for colour scheme used throughout all panels. Black datapoints denote extinct taxa. Shape denotes higher taxon; see respective legend for each panel.

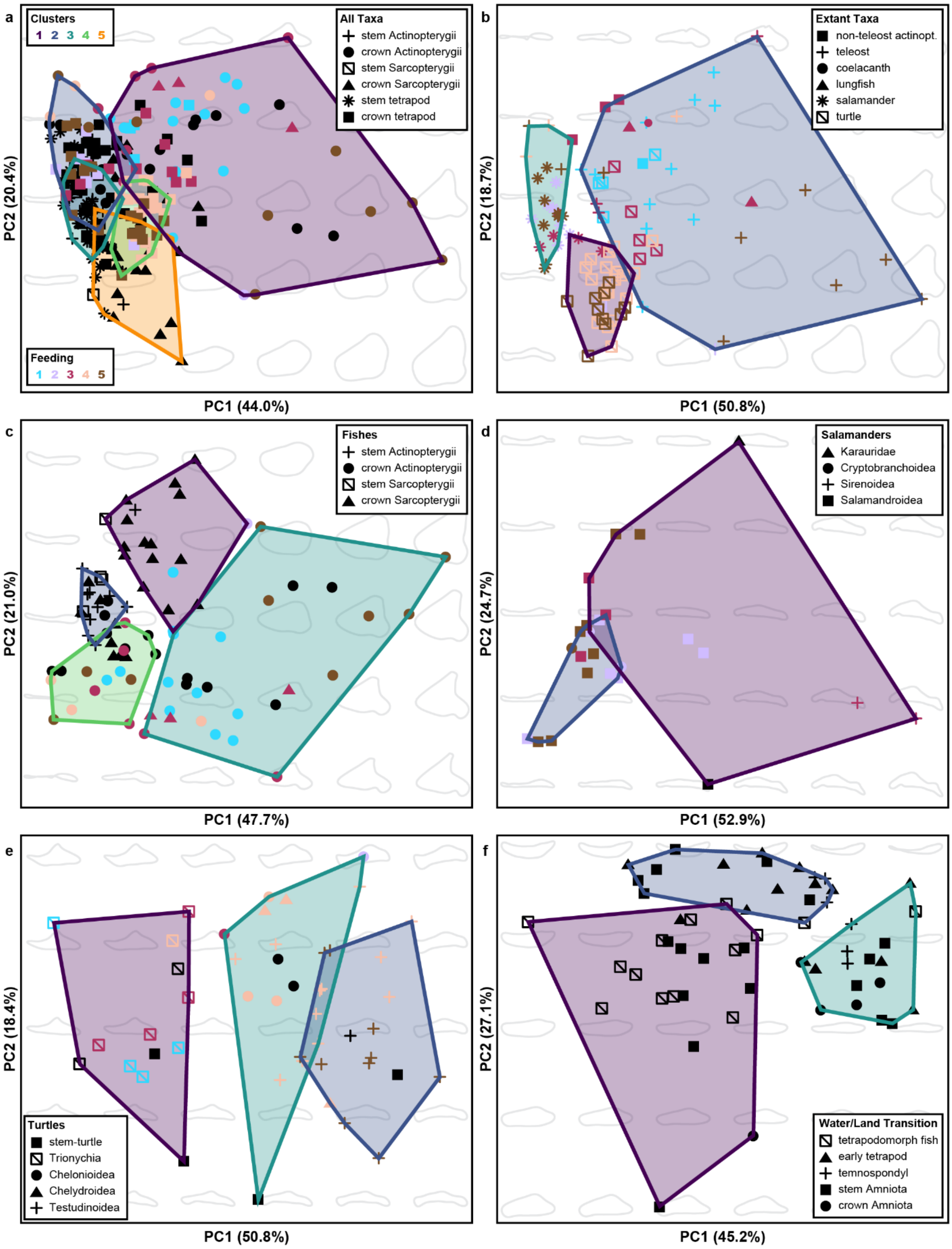
a
Clusters
1 2 3 4 5
All Taxa
stem Actinopterygii
crown Actinopterygii
stem Sarcopterygii
crown Sarcopterygii
stem tetrapod
crown tetrapod
Feeding
1 2 3 4 5
PC2 (20.4%)
PC1 (44.0%)
b
Extant Taxa
non-teleost actinopt.
teleost
coelacanth
lungfish
salamander
turtle
PC2 (18.7%)
PC1 (50.8%)
c
Fishes
stem Actinopterygii
crown Actinopterygii
stem Sarcopterygii
crown Sarcopterygii
PC2 (21.0%)
PC1 (47.7%)
d
Salamanders
Karauridae
Cryptobranchoidea
Sirenoidea
Salamandroidea
PC2 (24.7%)
PC1 (52.9%)
e
Turtles
stem-turtle
Trionychia
Chelonioidea
Chelydroidea
Testudinoidea
PC2 (18.4%)
PC1 (50.8%)
f
Water/Land Transition
tetrapodomorph fish
early tetrapod
temnospondyl
stem Amniota
crown Amniota
PC2 (27.1%)
PC1 (45.2%)

Figure S7. **Spectral clustering analysis of key taxon partitions, as distributed throughout morphospace, colour-coded using a 5-category feeding mode classification**. (**a**) All taxa (n = 214). (**b**) Extant taxa (n = 95). (**c**) Fishes (n = 84). (**d**) Salamanders (n = 24). (**e**) Turtles (n = 48). (**f**) Taxa spanning the tetrapod water-to-land transition (n = 58). Coloured polygons represent cluster assignments based on PC axes >5%. Coloured datapoints indicate feeding modes for extant taxa; see legend in (a) for colour scheme used throughout all panels. Black datapoints denote extinct taxa. Shape denotes higher taxon; see respective legend for each panel.

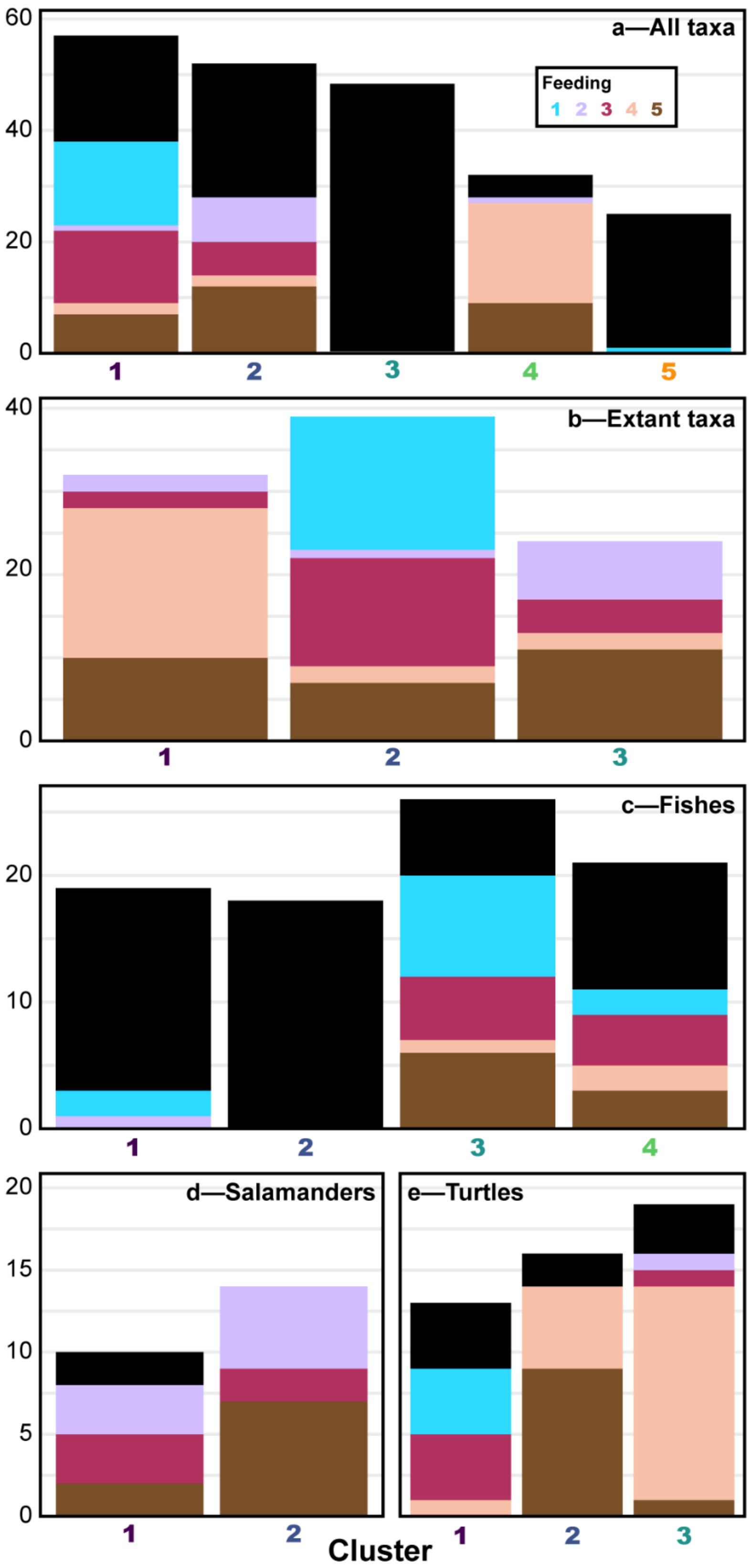
60
40
20
0
a—All taxa
Feeding
1 2 3 4 5
1
2
3
4
5
40
20
0
b—Extant taxa
1
2
3
20
10
0
c—Fishes
1
2
3
4
20
15
10
5
0
d—Salamanders
1
2
e—Turtles
1
2
3
Cluster

Figure S8. **Distribution of feeding modes across morphological clusters, obtained via spectral clustering analysis of key taxon partitions, colour-coded using a 5-category feeding mode classification.** (**a**) All taxa (n = 214). (**b**) Extant taxa (n = 95). (**c**) Fishes (n = 84). (**d**) Salamanders (n = 24). (**e**) Turtles (n = 48). Coloured bars indicate feeding modes for extant taxa; see legend in (a) for colour scheme used throughout all panels. Black denotes extinct taxa. Coloured labels along *x*-axis correspond to cluster polygons in Figure S7.

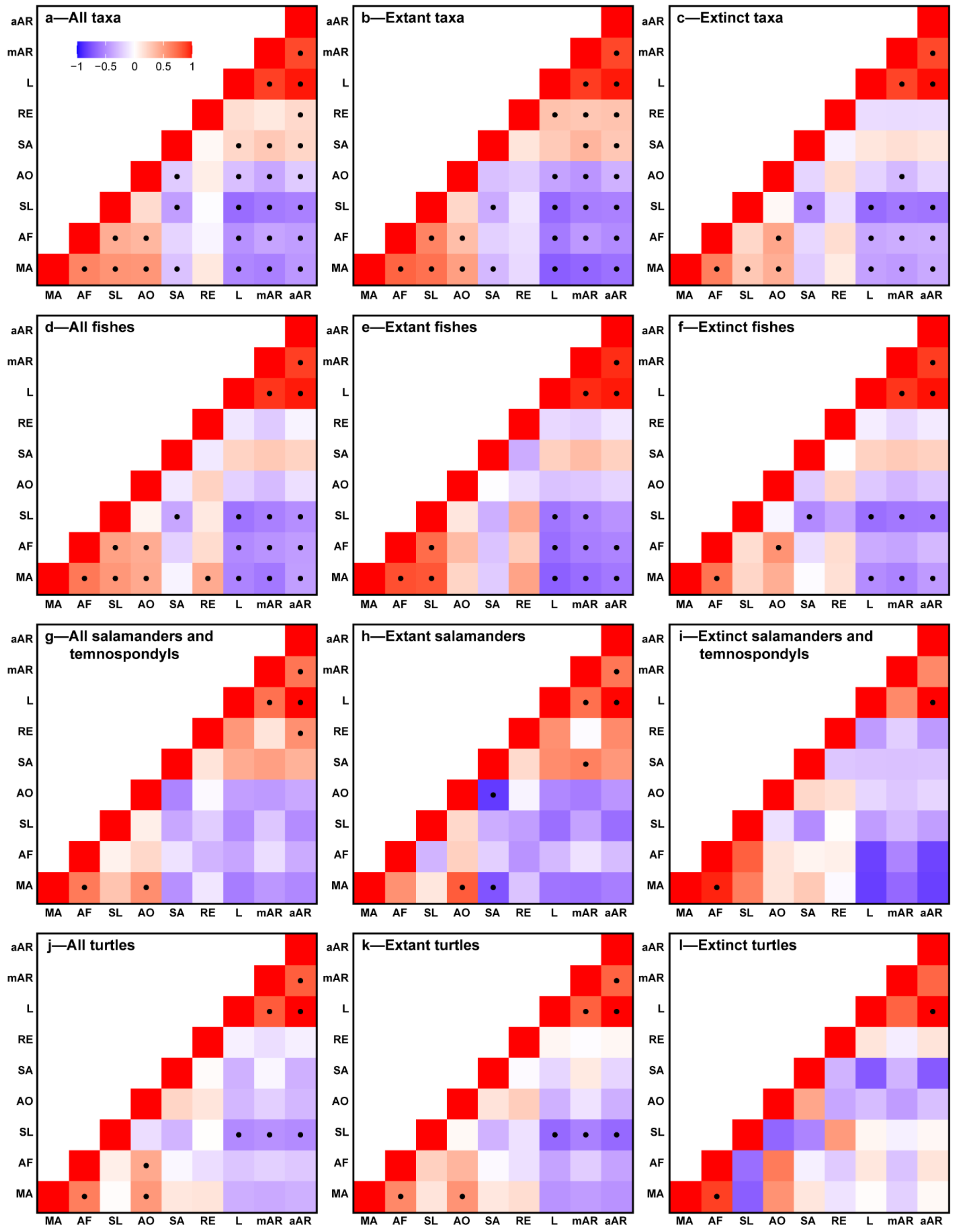
a—All taxa
b—Extant taxa
c—Extinct taxa
d—All fishes
e—Extant fishes
f—Extinct fishes
g—All salamanders and temnospondyls
h—Extant salamanders
i—Extinct salamanders and temnospondyls
j—All turtles
k—Extant turtles
l—Extinct turtles
−1 −0.5 0 0.5 1
MA AF SL AO SA RE L mAR aAR

Figure S9. **Heatmaps depicting pairwise correlation among functional metrics for key taxon partitions, including normalized jaw length**. (**a**) All taxa (n = 214). (**b**) Extant taxa (n = 95). (**c**) Extinct taxa (n = 119). (**d**) All fishes (n = 84). (**e**) Extant fishes (n = 34). (**f**) Extinct fishes (n = 50). (**g**) All salamanders and temnospondyls (n = 30). (**h**) Extant salamanders (n = 22). (**i**) Extinct salamanders and temnospondyls (n = 8). (**j**) All turtles (n = 48). (**k**) Extant turtles (n = 39). (**l**) Extinct turtles (n = 9). See scale bar in (a) for colour scale used throughout all panels. Cells marked with a dot are significant at $p<0.05$. See Methods for abbreviations of functional metrics.
